# Acute stress reshapes third-party punishment and help decisions: Behavioral evidence and neurocomputational mechanisms

**DOI:** 10.1101/2023.06.14.544887

**Authors:** Huagen Wang, Xiaoyan Wu, Jiahua Xu, Ruida Zhu, Sihui Zhang, Zhenhua Xu, Xiaoqin Mai, Chao Liu, Shaozheng Qin

## Abstract

People tend to intervene in others’ injustices by either punishing the transgressor or helping the victim. Injustice events often occur under stressful circumstances. However, how acute stress affects a third party’s intervention in injustice events remains open. Here, we show a stress-induced shift in third parties’ willingness to engage in help instead of punishment by acting on emotional salience and central-executive and theory-of-mind networks. Acute stress decreased the third party’s willingness to punish the violator and the severity of the punishment and increased their willingness to help the victim. Computational modeling revealed a shift in intervention severity bias from punishment toward help under stress. This finding is consistent with the increased dorsolateral prefrontal engagement observed with higher amygdala activity and greater connectivity with the ventromedial prefrontal cortex. A brain connectivity theory-of-mind network predicted stress-induced severity bias in punishment. Our findings suggest a neurocomputational mechanism of how acute stress reshapes third parties’ decisions by reallocating neural resources in emotional, executive and mentalizing networks to inhibit punishment bias and decrease punishment severity.

## Introduction

Humans are often willing to interfere in the injustices of others by either punishing the transgressor or helping the victim, even if they are not involved in the unfair event(Fehr et al., 2002). This extraordinary feature of human society allows us to establish, broadcast and enforce social norms (Fehr & Fischbacher, 2003, 2004). People typically prefer to punish the offender rather than help the victim in justice restoration (FeldmanHall et al., 2014; Stallen et al., 2018). Such preferences, however, are often context-dependent (David et al., 2017). Unpredictable and uncontrollable events are ubiquitous in daily life, and people frequently respond to norm violations under stressful situations, which is a reality that deserves special attention. Stress can profoundly impact psychological, neurophysiological, and brain networks involved in third-party decision-making, thereby influencing the affective response, motivation, and subjective value assigned to the decisions (Phelps et al., 2014; Ulrich-Lai & Herman, 2009).

The dual process theory of human decision-making emphasizes two distinct systems: system 1 is intuitive, fast, and regulated mainly by the limbic system, including the amygdala, whereas system 2 is reflective, slow, deliberate, and regulated mainly by the prefrontal cortex (Hallsson et al., 2018; Kahneman, 2013). Consistently, according to the biphasic-reciprocal model, stress-related hormones and neurotransmitters increase activity in the emotional salience network at the expense of the executive control network under acute stress(Hermans et al., 2014). Considerable evidence has shown that exposure to uncontrollable stress can shift brain function from a thoughtful, reflective mode toward a more rapid reflexive response regulated by higher-order prefrontal and primitive neural circuits, respectively(Arnsten, 2015; Phelps et al., 2014). Such stress-sensitive neural systems, at least in part, overlap with functional brain systems and networks involved in the decision-making process during third-party responses. Third-party decision-making under stress may be a “tend and befriend” pattern, meaning that decision behavior can be both adaptive and altruistic to promote survival and well-being at the individual and group levels(Taylor, 2006), and the impact of this stress on social decision-making is also supported at the neural level. Acute stress can lead to increased activation of “theory of mind” (ToM) brain regions (Tomova et al., 2017), which is fundamental for becoming more other-oriented with a preference for providing help (FeldmanHall et al., 2015).

Third-party punishment and help are both altruistic and fundamentally share a common neuronal mechanism with some distinct processes (Stallen et al., 2018). When facing injustice, both third-party responses correlate with the activation of fairness violations, e.g., the anterior insula (AI), anterior cingulate cortex (ACC), dorsolateral prefrontal cortex (DLPFC), striatal reward circuitry and emotional regulation systems, including the amygdala and ventromedial prefrontal cortex (vmPFC)(Stallen et al., 2018; Xie et al., 2022). Enhanced ventral striatal activity is further associated with punishing an offender instead of helping a victim (Stallen et al., 2018). On the other hand, the temporoparietal junction (TPJ), precuneus, and mid-temporal regions are more associated with an individual preference for help rather than punishment(Civai et al., 2019). In other circumstances, activity in the participants’ right dorsolateral prefrontal cortex (rDLPFC) increased more after experiencing advantageous unfairness and engaging in help instead of punishment, whereas those who punish more after experiencing disadvantageous unfairness demonstrated significantly decreased activity in the rDLPFC(Xie et al., 2022). Although studies have demonstrated that stressed individuals have greater altruistic (greater trust, sharing, and generosity)(Takahashi et al., 2007; von Dawans et al., 2012a), cooperative, and other-oriented tendencies (Nickels et al., 2017a), the neurocognitive mechanisms underlying the trade-off between third-party help and punishment under acute stress remain unclear.

In addition to the trade-off of third-party help and punishment at a behavioral level, it remains unclear whether the two candidate decisions involve differential neurocomputational processes; for instance, whether and with what severity to intervene in an injustice event under uncontrollable stress(Hu et al., 2015; Stallen et al., 2018). The computational model allows us to decipher signals that are not expressed by the behavioral indicators, e.g., in value utility calculations, people rely only on intuition and habit in the allocation of some choices (e.g., helping), while others (e.g., punishing) are considered more carefully and deliberately. We can identify specific neural circuits linked to specific decision variables and processes by combining economic computational tools and neuroimaging approaches. Therefore, it is critical to investigate the neurocomputational bases of third-party intervention under acute stress by integrating economic paradigms, computational tools, and neuroimaging approaches. Based on the biphasic-reciprocal model of the stress-induced transition from deliberate to intuitive under stress, we hypothesized that people under stress would prefer helping when third-party punishment and helping are in conflict because punishment is more executive control–dependent. We expected that acute stress would decrease behaviors that require adequate cognitive control while increasing behaviors that recruit habitual and reflexive regulation. According to empirical evidence from many previous studies, helping victims mainly reflects more habitual behavior in stressful situations, which aligns with “tend and befriend” due to its social approach property (Buchanan & Preston, 2014; Taylor, 2006). However, the willingness to punish requires greater DLPFC executive function (Buckholtz et al., 2008).

To test the above hypotheses, we set up an event-related fMRI study in conjunction with an economic paradigm of third-party intervention and computational modeling approach to examine the behavioral and neurocomputational bases of third-party interventions under acute stress. A classic cold pressor test (CPT) was used to induce acute stress (**Fig 1A**), and a novel third-party intervention game (TPIG) (**Fig 1E&1F**) was designed to detect the willingness/preference and severity of third-party decisions. The two subcomponents were embedded in the decision and transfer phases, respectively. During the decision phase, participants decided whether to intervene in injustice events related to others. In the subsequent transfer phase, participants decided how many tokens to use to punish or help others if they chose to deduct the proposer or add the receiver, respectively. We constructed a computational model based on the assumption that participants made decisions by integrating self-payoff and other-regarding inequality aversion. This model allowed us to estimate the parameter values reflecting individual degrees of punishment and help, with a greater punishment severity or increased help requiring more tokens to diminish the violator’s advantageous status or promote the victim’s disadvantageous status, respectively. We expected to observe neural changes in brain functioning related to the theory of mind (e.g., ACC and TPJ) and subjective value computations (e.g., vmPFC and PCC)(Bartra et al., 2013) under stress because deciding how seriously to punish or how extensively to help is strongly considered in relation to others (i.e., payoff and feelings) and the value calculation of selectable options.

**Fig 1.**
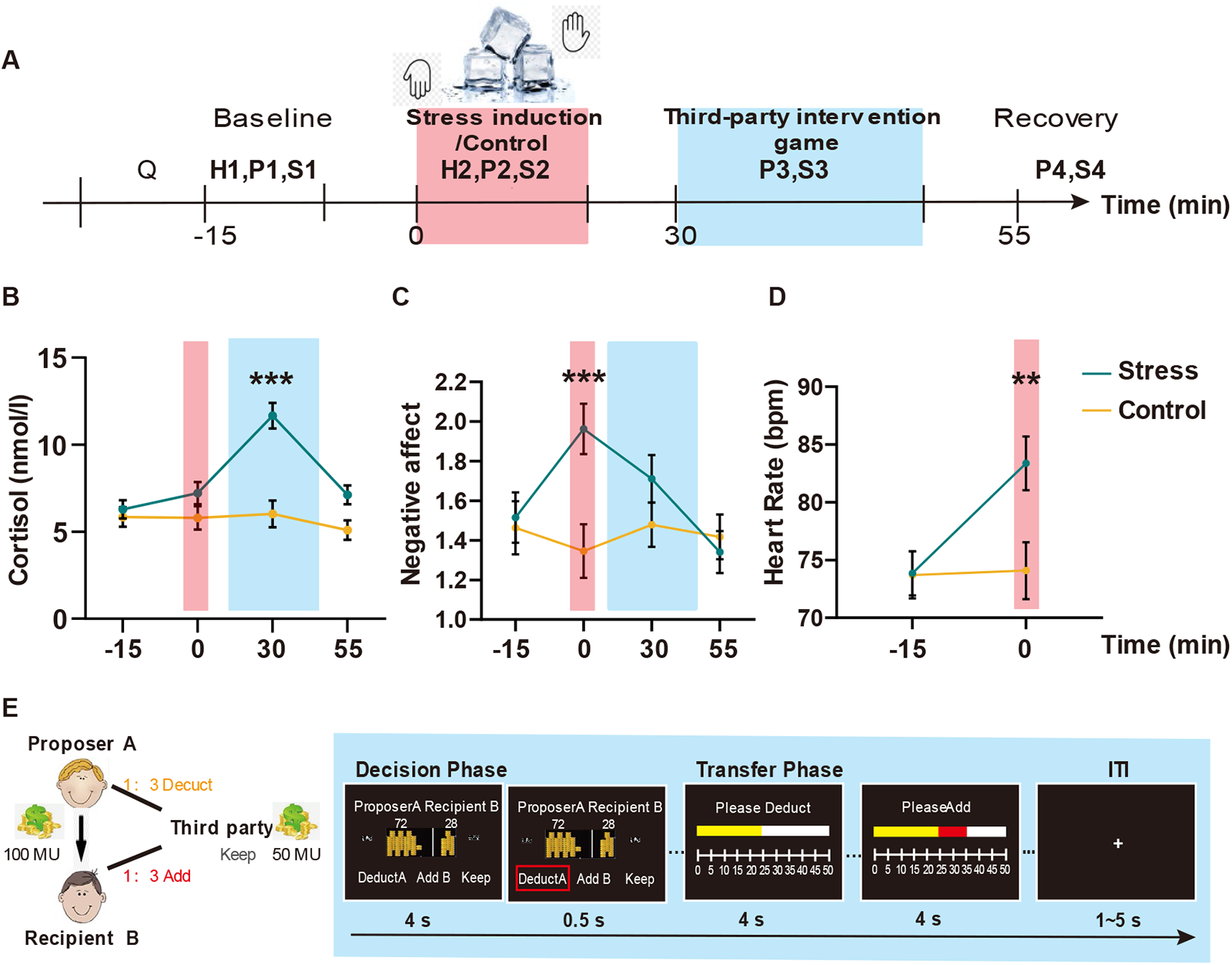
Experimental procedure, stress induction and third-party intervention game. **(A)** Timeline depicting the experimental procedure. Four saliva samples with concurrent subjective affect ratings were collected throughout the task (S1-S4 and P1-P4). Heart rate was collected during stress induction (H1) and the third-party intervention game (H2). **(B-D)** The curves depict salivary cortisol concentration, negative affect, and heart rate in the control and stress groups. The red and blue boxes represent the CPT and TPI time windows. **(E)** An illustration of the third-party intervention game with a sample trial. In each trial, participants were asked to choose from three options within 4 s (i.e., subtract “A”, add “B”, or keep) in the decision window, and they were instructed to decide how many MUs to use to reduce the MUs of A or increase the MUs of B within 4 s in the transfer window. If the participants chose to keep the money for themselves, the transfer phase windows were shown to the participants, but they were asked to select the zero option (meaning no tokens were used to intervene in other-regarding events). A 1–5 s intertrial interval was set between each window to dissociate the neural signal from each phase. \*\*\**P* <.001; \*\**P* <.01; error bars represent SEM.

## Results

### Acute stress induction with psychological, physiological, and endocrinal measures

We first examined the effectiveness of CPT-induced acute stress by analyzing salivary cortisol, heart rate and self-reported affect feelings in the stress and control groups. We conducted two-way analyses of variance (ANOVA) of Group (between-subject factor: stress vs. control) × Time points (within-subject factor) on salivary cortisol level, negative affect, and heart rate. This analysis revealed a significant Group × Time interaction for salivary cortisol level (*F*(2.22, 110.75) = 11.231, *P* < 0.001, η_p_^2^ = 0.183; *P* < 0.001), self-reported negative affect (*F*(2, 98) = 7.38, *P* < 0.001, *η*_p_^2^ = 0.131; *P* < 0.01) and heart rate (*F* (1, 49) = 18.19, *P* < 0.001, *η*_p_^2^ = 0.271). Bonferroni-corrected post hoc t tests revealed that during CPT, the stress group exhibited higher levels of salivary cortisol (P< 0.001, **Fig 1B**) and negative feelings (P< 0.001, **Fig 1C**) than the controls and also exhibited elevated heart rates (P< 0.01, **Fig 1D**). These results indicate that CPT induced a prominent acute stress response, aligning with previous studies’ findings using a similar stress paradigm. We compared relevant individual personality (e.g., perceived stress, empathetic concern) and endocrine variables (i.e., baseline basal testosterone and oxytocin level) between groups to exclude other possible individual effects that might confound the acute stress effect. These comparisons revealed no significant differences in these measures (all P > 0.131; see **Supplementary Table 1**). Thus, any observed differences in behavioral and brain responses between groups could be attributed to acute stress but not personality and baseline endocrine levels.

### Acute stress affects third-party decisions between punishment and help in response to inequality events

We next investigated the effect of acute stress on behavioral performance in the third-party intervention game (TPIG). We conducted a two-way ANOVA of Group (between-subject factor: stress vs. control) × Fairness (within-subject factor: 50:50 vs. 60:40 vs. 70:30 vs. 80:20 vs. 90:10) × Intervention type (within-subject factor: punish vs. help) regarding the likelihood of each choice in the decision phase. This analysis revealed a trend of Group × Fairness × Intervention interaction (*F* (3, 150) = 2.811, *P* = 0.108, *η*_p_^2^ = 0.040). A further exploratory analysis revealed that acute stress affects decision-making only in extremely unfair conditions (i.e., 80:20 and 90:10 trials), with an increase in third-party decisions to help (P< 0.05 in 80:20 and 90:10 conditions) but a decrease in third-party decisions to punish (*P* < 0.05 in 80:20 and 90:10 conditions) in the stress group compared with the control group (**Fig 2A**). The parallel analysis of third-party intervention severity data revealed a similar pattern only in the extremely unfair condition. That is, acute stress decreased the third party’s willingness to punish the norm violator (*P* < 0.01 in the 80:20 condition and P< 0.05 in the 90:10 condition) but had no effect on helping the receiver (*P* > 0.05) (**Fig 2B**). We restricted our further analyses to focus on the stress-by-intervention type in highly unfair conditions.

We conducted two-way ANOVA of Group × Intervention type on the choice rate of the decision phase of extremely unfair trials and found a significant interaction (*F* (1, 50) = 4.033, *P* = 0.05, *η*_p_^2^ = 0.075). Bonferroni-corrected post hoc t tests revealed that stress increased the likelihood of choosing helping (*t _help_* (50) = 2.11, *P* < 0.05, 95% CI: 0.01, 0.31) but slightly decreased the likelihood of choosing punishment compared with the control group (*t _punishment_* (50) = -1.94, *P* = 0.058, 95% CI: -0.30, 0.01) (**Fig 2C**). Using the same two-way ANOVA on the contribution in the transfer phase (*F* (1, 50) = 3.13, *P* = 0.08, *η*_p_^2^ = 0.059), we found that stress decreased the contribution made to punish the norm violator (*t* _help_ (50) = 0.79, *P* = 0.44, 95% CI: -2.06 to 4.71; *t* _p*unishment*_ (50) = -2.08, *P* < 0.05, 95% CI: -6.01, -0.10) but had no effect on the contribution made to help the receiver (**Fig 2D**). These results indicate that acute stress when facing other extreme injustice events decreases third parties’ willingness to punish norm violators and the severity of the punishment but increases their willingness to help victims.

### Acute stress alters the latent computations of third-party intervention behaviors

To further investigate the latent cognitive computations underlying how acute stress modulates third-party intervention behaviors, we constructed four plausible computational models (see methods) to fit participants’ behaviors in the transfer phase to model the mental computations of punishment severity and the extent of help. As shown in **Fig 2E**, the model comparisons revealed that our behavioral data could be best fitted by the other-regarding inequality aversion model. This model allows us to separate the latent computations regarding the extent of help (i.e., how averse a person is to observe someone else being hurt) and the severity of punishment (i.e., how averse a person is to observe someone else hurt others), which are quantified by parameters *β* and *α,* respectively. We computed the value of parameter α minus parameter *β* as an index of punishment bias, ranging from -1 to 1. A larger value indicates that individuals tend to punish the norm violator more severely than the extent to which they help the victim. The one sample t test revealed that acute stress reduced the punishment bias compared to that demonstrated by the controls (**Fig 2F**, *t* (50) = -2.62, *P* < 0.05, 95% CI: -0.50, -0.07). We further examined the association between model-based parameters and model-free behavior outputs to confirm the psychological meaning of the punishment bias value. We found a positive correlation of the punishment bias value with the relative punishment rate in the decision phase (i.e., the frequency of choosing punishment minus that of choosing helping, **Fig 2G**, r = 0.745, *P* < 0.01) as well as with the relative punishment severity in the transfer phase (i.e., the contribution donated to punishment minus the contribution donated to helping, see **Fig 2H**, r = 0.792, *P* < 0.01). These results indicate that acute stress reduces punishment bias by decreasing punishment severity but increasing the extent of help.

**Fig 2.**
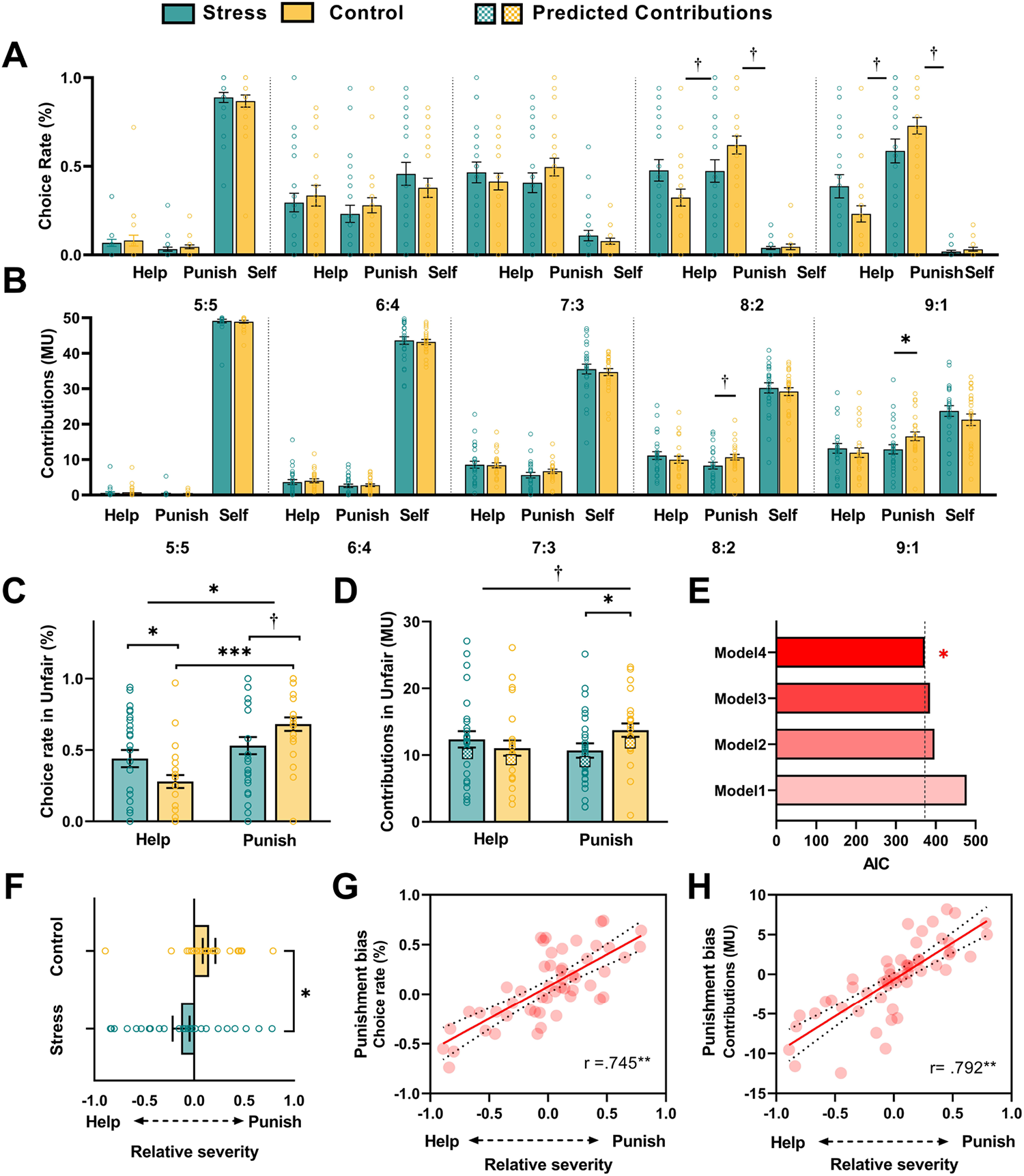
Behavioral measures and computational modeling of the third-party intervention game. (**A&C**) Stress decreased the punishment rate but increased the help rate in the unfair condition (80:20 & 90:10). (**B&D**) In the unfair condition, stress decreased the severity of punishment. The scatter plot depicts the correlation between the actual and predicted contributions based on computational modeling. (**E**) An overview of model comparisons: Models 1 and 2 represent the baseline model and the inequality aversion model. Model 3 represents the other-regarding inequality aversion model with a shared parameter of relative severity preference. Model 4 represents the other-regarding inequality aversion model with two parameters of the severity of punishment and help separately. (**F**) The difference between the stress and control groups in the relative severity preference based on the computational model (*α* – *β*). (**G**) The actual choice rate differences between punishment and help (punishment bias) in all conditions as a function of the model-estimated relative severity. (**H**) The actual contribution differences between punishment and help (punishment bias) in all conditions as a function of the model-estimated relative severity. Each dot represents the data of a single participant. \*\*\**P* <.001; \*\**P* <.01; \**P* <.05; †*P* <.01. Error bars represent the SEM.

### Stress increases amygdala and prefrontal network integration in third-party intervention responses to inequality events in the decision phase

To investigate the overall effect of acute stress on brain systems involved in the third-party intervention game (TPIG), we conducted a GLM with a parametric design(Büchel et al., 1998) to identify brain regions coding “Inequity” on the decision phase and the overall effect induced by stress. First, we found higher activation in the left amygdala and the bilateral insula (height threshold *P* < 0.001; extent threshold *P* _FWE_ < 0.05, small-volume corrected for the anatomically defined amygdala mask) and TPJ (**Table S2**, height threshold *P* < 0.001; extent threshold *P* _FWE_ < 0.05, whole-brain corrected) in the stress group than in the control group. These results confirm earlier findings indicating the essential role of the emotional salience network in the acute stress response, indicating the validity of CPT-induced changes in brain functioning. Critically, we found that the activity in the rTPJ, VLPFC, and DLPFC was positively correlated with distributional inequity between the proposer and the recipient for both the stress and control groups (Table S3, height threshold of P < 0.001; extent threshold P _FWE_ < 0.05, whole-brain corrected). This finding aligns with the meta-analytical result of neural processing of inequity (Feng et al., 2015). In addition, we found that the correlation between the right amygdala and the degree of distributional inequity was significantly higher in control individuals than in stressed individuals (**Fig 3A** & Table S3).

To further investigate the effects of acute stress on the amygdala-centric emotional salience network, we conducted a whole-brain psychophysiology interaction (PPI) analysis using the amygdala as a seed of interest. This analysis revealed higher functional connectivity of the right amygdala with the vmPFC in response to trials where participants chose the punishment option in the stress group but not in the control group (**Fig 3A&B, Table S5**, height threshold *P* < 0.001; extent threshold of *P* _FWE_ < 0.05, whole-brain corrected). Using a mediation model, we further found that acute stress reduced the punishment rate by increasing amygdala-vmPFC communication (**Fig 3C**). This result indicates that although stress decreases the insensitivity of the right amygdala to the degree of inequity, it fundamentally changes the communication between the amygdala and the emotion regulation network and further decreases the frequency of choosing the punishment option in the decision phase.

Moreover, we compared brain activation maps for trials in which punishment was selected with trials in which help was selected between the stress and control groups. In the control group, we found higher activity in the DLPFC when selecting the helping option than when the punishment option was selected (**Fig 3D**). However, the stress group showed an inverse pattern: higher activity in the DLPFC was observed when selecting the punishment option than when selecting the helping option (**Fig 3D**, initial threshold *P* < 0.001; extent threshold of *P* _FWE_ < 0.05, whole-brain corrected). We also compared the punishment and help choices separately between the stress and control groups. This contrast revealed that only during the punishment choices did acute stress induce higher activity in the right DLPFC, right TPJ (rTPJ), and right PCC (rPCC) (**Fig 3E-G**, initial threshold *P* < 0.001; extent threshold *P* _FWE_ < 0.05, whole-brain corrected). These results indicate that punishment activates the DLPFC, rTPJ, and rPCC to a greater extent in the stress group than in the control group. However, the parallel analysis of brain activity and connectivity associated with the help trials revealed no clear effects when comparing the acute stress and control groups. Together, these results indicate that punishment, but not helping, might be a more intuitive and automated process, since we observed that the control group was more likely to choose the punishment option and exerted less cognitive effort to make punishment decisions.

**Fig 3.**
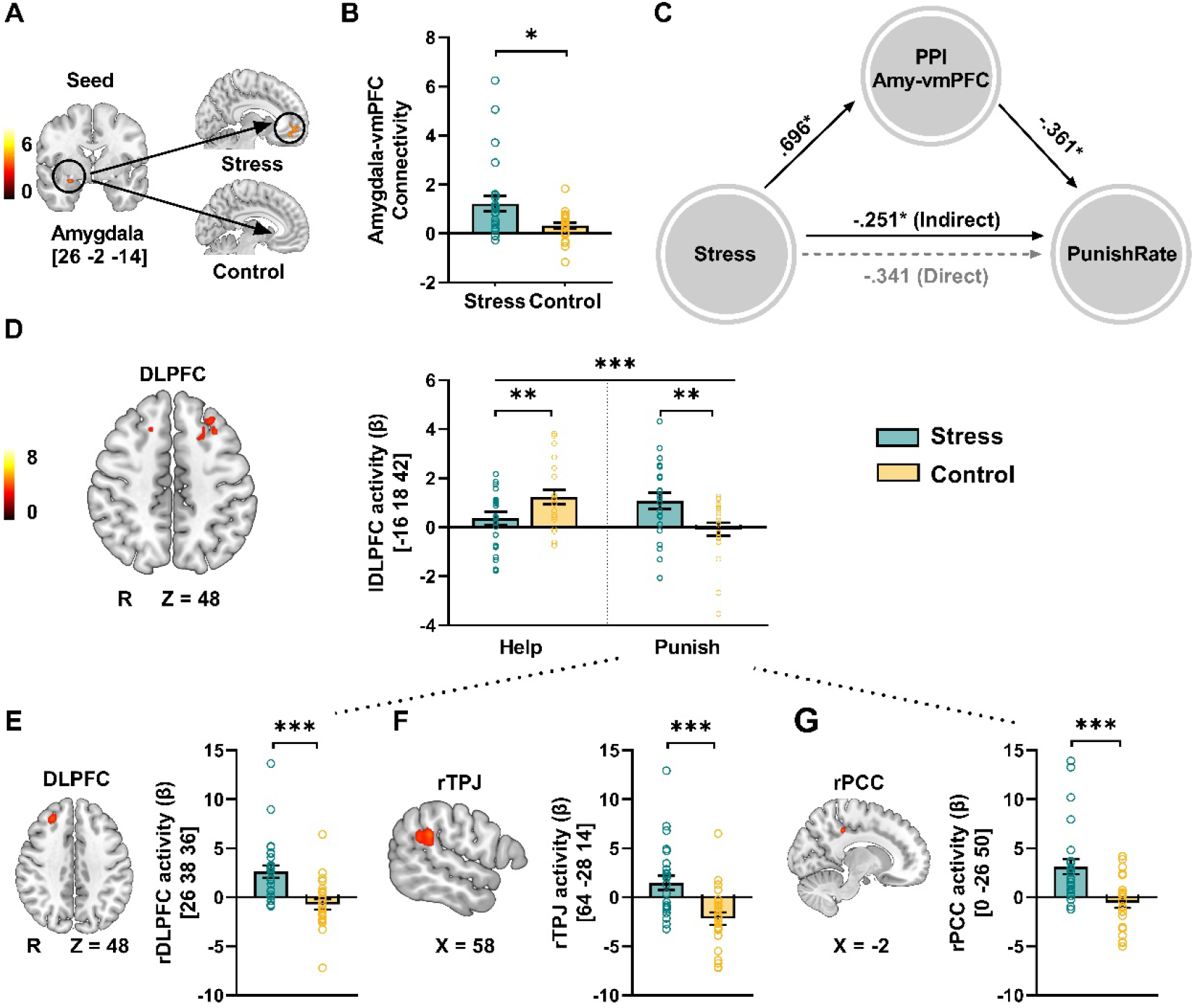
Stress-induced changes in amygdala emotional salience and prefrontal executive networks in the decision phase. **(A&C)** Stress-induced differences in brain representation of the degree of inequality and the functional connectivity between the amygdala and vmPFC. **(A)** The correlation between the degree of inequity and the activation of the right amygdala was significantly higher in the control group than in the stress group (small volume correction (SVC), cluster corrected at *P* _FWE_ < 0.05, following an initial threshold of *P* < 0.001, **Table S3**). **(B)** PPI analysis based on the results of (A) with the right amygdala as a seed showed that the stress group had increased functional connectivity of the right amygdala and the vmPFC during the decision stage in trials in which participants selected the punishment option. \**P* < 0.05; Error bars represent the SEM. **(C)** The mediating effect of amygdala-vmPFC connectivity on the association between acute stress and the punishment rate (i.e., the frequency of selecting the punishment option in the decision phase). **(D&G)** Stress-induced neural activity in the decision phase. **(D)** Relative to the control group, the stress group demonstrated stronger DLPFC activation when selecting the punishment option than when selecting the help option (initial whole-brain threshold *P* < 0.001, cluster corrected *P* _FWE_ < 0.05 for left DLPFC). **(E&G)** Relative to the control group, the stress group had stronger activation in the rDLPFC, rTPJ and rPCC in trials in which participants selected the punishment option (initial threshold *P* < 0.001; cluster corrected *P* _FWE_ < 0.05), and the effect was not significant in trials in which participants selected the help option.

### Acute stress alters brain functional connectivity and latent computations in the transfer phase

To investigate the neurocomputational mechanisms of how acute stress alters functional brain systems involved in third-party intervention responses to inequality events, we first conducted a parametric modulation analysis with trial-by-trial subjective values (the “Utility” in the computational model) as a parametric modulator during the transfer phase. This analysis identified that the vmPFC, PCC, and several other regions were critical for the integrated subjective value of the transfer magnitude by maximizing utility for both stressed and control groups (**Fig 4B**; **Table S6**, height threshold P < 0.001, extent threshold P _FWE_ < 0.05 whole-brain corrected). Interestingly, after correcting for multiple comparisons, we did not observe consistent differences in brain regions in subjective value representation between the stress and control groups.

Motivated by the behavioral effect on the stress-induced decrease in prosocial punishment in the transfer phase, we investigated the neurocomputational correlates of punishment severity bias (punishment relative to help, *α-β*) during the transfer stage. Whole-brain multiple regression analysis revealed that punishment severity bias was more strongly correlated with the activity of the ACC (a height threshold P < 0.001, an extent threshold P_FWE_ < 0.05 SVC corrected), PCC, and rTPJ in stressed versus control participants (height threshold P < 0.001, P_FWE_ < 0.05 whole brain corrected). Specifically, in the stress group, we observed a positive correlation between punishment bias and activity in the ACC, PCC, and rTPJ, while the control group showed negative correlations (**Fig 4A**).

We further conducted a moderated mediation model to confirm that acute stress moderated the pathway from neural activation to behavioral outputs. In this model, the activation of the ACC (or PCC or rTPJ) in the transfer stage was used as an independent variable, punishment severity bias (*α-β*) was used as a mediator, stress manipulation (stress or control) was used as a moderator, and punishment contribution bias (punishment contribution minus help contribution) was used as a dependent variable. This analysis revealed that acute stress acts as a moderator variable in a mediation model that showed that the activation of the ACC and PCC affects behavioral performance through its effect on punishment bias (**Fig 4C**). Specifically, the stressed participants must recruit more ACC and PCC cognitive resources for a larger punishment severity bias and impose a higher punishment contribution. In contrast, individuals who did not experience acute stress imposed a higher punishment contribution, as they showed a larger punishment severity bias with lower ACC and PCC resource consumption. These results indicate that acute stress moderates the mediatory role of latent computations on the association between neural activity and behavior outputs.

**Fig 4.**
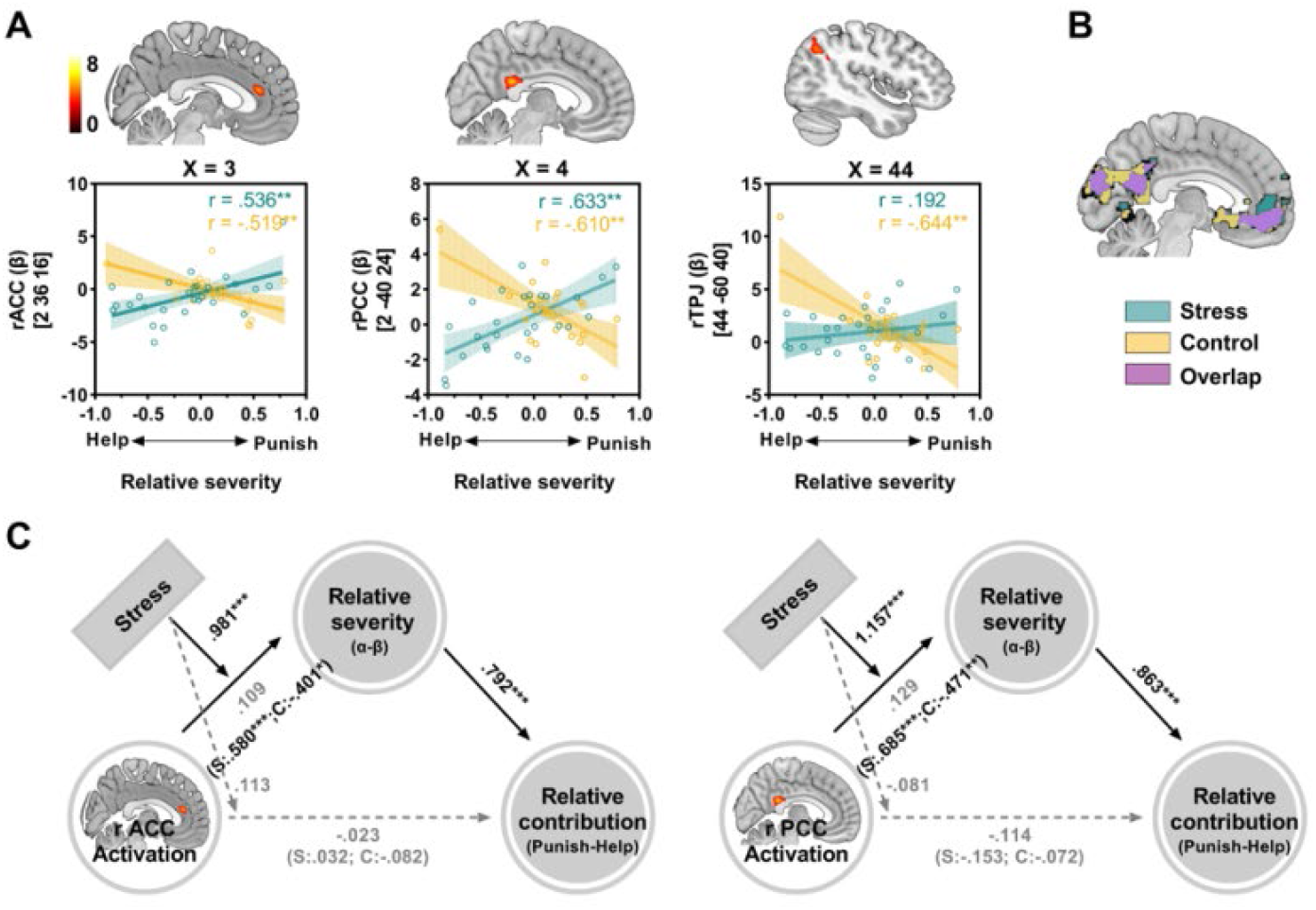
Stress-induced changes in brain-behavior associations with severity computations in the transfer stage. (**A**) Brain-behavior relationships between the stress-induced shift of punishment bias (*α-β*) and activation in the rACC, PCC and rTPJ. Scatter plots depict the correlations between punishment bias (*α-β*) and activation in these regions. (**B**) Parametric modulation with trialwise Utility revealed a significant correlation with BOLD responses in the vmPFC and PCC for both the stress and control groups. (**C**) The moderated mediation models depict that the punishment bias (*α-β*) could account for the indirect associations of value computation in the activity of the rACC (left panel) and rPCC (right panel) with the punishment severity bias (punishment contribution minus help contribution). Acute stress moderated the correlation between value computation and punishment bias. All significant clusters were determined by a height threshold of *P* < 0.001 and an extent threshold *P* _FWE_ < 0.05. SVC.

## Discussion

This study investigated the behavioral and neurocomputational substrates of how acute stress impacts a third party’s intervention in response to injustice events. Behaviorally, acute stress led to a decrease in the third-party’s willingness to punish the violator and the severity of the punishment but an increase in their willingness to help the victim. Computational modeling further revealed a shift in intervention severity bias from punishment to help under acute stress. Critically, we further found that acute stress induced higher activity in the insula, amygdala, and TPJ, reflecting a hyperactive state of the emotional salience network. This is consistent with greater connectivity with the vmPFC and higher dorsolateral prefrontal engagement. Brain connectivity in the theory-of-mind network positively predicted the stress-induced punishment severity bias. These findings will be discussed in a framework of behavioral and neurocomputational mechanisms of how acute stress affects third parties’ decisions through acting emotional salience, prefrontal control and mentalizing networks.

On a behavioral level, our findings support the “tend-and-befriend” theory, and a growing body of studies has demonstrated that stressed individuals have greater altruistic (more trust, sharing, and generosity) (Takahashi et al., 2007; von Dawans et al., 2012b), cooperative, and other-oriented tendencies(Nickels et al., 2017b; Youssef et al., 2018). Evolutionally, an appropriate and modulated stress response is the core of survival, and a “tend-and-befriend” strategy can lead to a good reputation and promote self and offspring safety(Taylor, 2006). In contrast, punishment entails a cost for both the punisher and the punished, and it is expensive and inefficient. The threat of retaliation and vengeance from the target might lead individuals to avoid punishment when other nonconfrontational options are available, especially in uncontrollable stress situations(Dreber et al., 2008; Rockenbach & Milinski, 2006). From the perspective of reciprocity, helping an individual in need might create opportunities for direct reciprocity from that individual or an uninvolved bystander(Raihani & Bshary, 2015; Sylwester & Roberts, 2013). This aligns with the notion that acute stress may trigger a social approach, motivating individuals to build social bonding with other social members.

By leveraging computational modeling, we modeled the process by which participants integrated value utility and assigned different mental weights and subjective biases to the allocation of amounts to punishing the offender and helping the victim when making a norm recovery. The model parameter showed that stressed participants have smaller punishment bias. That is, when subjective value utility calculations are made in stressful situations, people devote more of their psychological preferences to deciding how to help the victim than how to punish the offender. These results from the computational model further suggest that helping, rather than punishment, is the more intuitive and efficient response pattern, both in terms of deciding whether to punish or help and in deciding the magnitude of punishment and helping.

On a neuroimaging level, we found that stressed individuals demonstrated increased DLPFC activity when selecting punishment options instead of help options, which is commonly associated with top-down executive function in controlling selfishness-related impulses (Sanfey et al., 2003), social norm compliance in adjusting inequity aversion (Knoch et al., 2006), and integrating distinct information streams for appropriate decisions(Zinchenko & Klucharev, 2017). Particularly, we found that stressed individuals demonstrate greater DLPFC, PCC, and rTPJ activation in those punishment options, and no consistent region is affected by stress in helping options. These results are consistent with the findings of previous studies showing that third-party punishment recruits the DLPFC critical for executive functioning by integrating multiple information streams for appropriate decisions(Zinchenko & Klucharev, 2017) (Sanfey et al., 2003). Research has shown that the PCC plays an important role in value-based decision-making, especially in terms of behavioral control and the evaluation of affordances (Grueschow et al., 2015; Pearson et al., 2011). Numerous studies have also linked neural activity in the rTPJ and PCC with mentalizing-related computations(Schurz et al., 2014) and strategic choice(Hill et al., 2017). Based on the biphasic-reciprocal model in which behavior patterns transition from deliberate to intuitive under stress, these results suggest that punishment in stressful situations reflects a relatively more complex aspect, which not only requires one to activate the mentalization function to consider the emotional state of the person in the situation but also requires investing cognitive resources in value calculation to weigh the pros and cons. In contrast, helping behavior under stress is more intuitive and straightforward and becomes a more adaptive and habitual decision with relatively lower cognitive and computational expenditure. Broadly, people typically prefer to punish the offender than help the victim for justice restoration (FeldmanHall et al., 2014; Stallen et al., 2018). In contrast, this considerate response was dramatically changed under stress, and punishment is no longer an optimal strategy in dealing with other-regarding injustice events since it requires more cognitive and mentalizing resource involvement; only then can people inhibit automated responses and make more prudent and adaptive decisions(Tomova et al., 2017).

In conjunction with local activation, we found that stress increased amygdala-vmPFC functional connectivity, and this connectivity, in turn, inhibited the punishment option, correspo to the early findings that the degree of inequity aversion is predictable from amygdala activity(Haruno & Frith, 2010). This result is consistent with previous studies that demonstrated that psychological stress and early life stress increased vmPFC functional connectivity with the amygdala(Arnsten, 2015; Carmichael & Lockhart, 2018; Ginty et al., 2019; Sinha et al., 2016). Simultaneously, animal research and human studies have suggested that top-down regulation of the vmPFC to the amygdala is implicated in emotion regulation and extinction learning(Burghy et al., 2013; Phelps & LeDoux, 2005). Overall, acute stress decreased the initial flexibility of the amygdala response to inequity, and stressed participants recruited more emotion regulation of the vmPFC to curb inequity aversion when making punishment decisions.

Furthermore, we found that the vmPFC, PCC, and several other regions represented the subjective utility value in the transfer stage in both the stress and control groups. This finding is consistent with the results of a large body of studies in that acute stress did not fundamentally change the subjective value computation circuits (Maier et al., 2015; Philiastides et al., 2010). We also found that stressed individuals demonstrated stronger rACC, rPCC, and rTPJ activities when computing the relative severity of punishment against help than those in the control group. The ACC has been shown to balance the motivational conflict between the immediate emotional reaction and maximize profit in the decision-making context (Botvinick, 2007; Sanfey et al., 2003). Moreover, we found that acute stress plays a moderating role in the direction of value computation of the rACC and rPCC to the relative severity preference between punishment and help and subsequently affects the transfer level of punishment relative to help. Specifically, the stressed participants needed to use more cognitive rACC and rPCC resources for a larger punishment severity bias and to impose a greater punishment contribution. In contrast, individuals in the control group only need to invest a low level of rACC and rPCC resource consumption in deciding how severely to punish. These findings further support the conclusion that helping rather than punishment is a more intuitive and efficient response pattern. Given that individuals under stress need to consume more energy precisely involved in calculation when making punishment decisions, to survive in the future, individuals under stress will reduce cognitive control and computational recruitment. Notably, since we only collected male samples, the results are more generalizable to men than women. Future research with a larger sample size is needed to determine possible sex differences under stress.

In conclusion, our study demonstrates a stress-induced shift in third parties’ tendency from punishment toward help by acting on emotional salience, central-executive and theory-of-mind networks, characterized by higher amygdala activity and greater connectivity with the vmPFC, an increase in dorsolateral prefrontal engagement, and connectivity of the mentalizing network in relation to the stress-induced severity bias of punishment. Our findings suggest a neurocomputational mechanism of how acute stress reshapes third parties’ decisions by reallocating value computations and neural resources in emotional, executive and mentalizing networks to inhibit punishment bias.

## Materials and Methods

### Participants

Fifty-three male volunteers were recruited and randomly assigned to either the stress condition (CPT) or the control condition; one participant was removed from the analyses, as he did not understand the task. The final sample included 52 participants (stress: n = 27). Participants who met the following conditions were excluded: hormonal contraception, prescription or drug consumption, smoking, alcohol abuse, a history of chronic disease or mental condition, and a major evaluation within two weeks. Furthermore, participants were instructed to abstain from physical activity, meals, and caffeine consumption for 2 hours prior to the study. All study procedures were approved by the Institutional Review Board of the State Key Laboratory of Cognitive Neuroscience and Learning at Beijing Normal University. All participants signed written informed consent.

### Experimental procedures

Testing sessions took place in the afternoon (1:30–5:30) to control variability in diurnal cortisol secretion. Upon arrival at the lab, participants completed questionnaires (psychological tests, see SI Appendix) for 20 minutes. Subsequently, the baseline heart rate (HR1) was recorded for 3 minutes, and the saliva sample (S1) and the Positive and Negative Affect Scale (PANAS, PA1, and NA1) were measured. Then, the participants were randomly assigned to CPT or control conditions. Heart rate (HR2) was recorded across the whole CPT for 3 minutes. The saliva sample (S2) and the PANAS (PA2 and NA2) were collected immediately after the CPT or a control manipulation.

Then, after 10 minutes of rest and 10 minutes of T1 structural image collection in the MRI scanner, the subjects completed the first run of the third-party intervention task (TPI). The third saliva sample (S3) and the PANAS (PA3 and NA3) were measured 25–30 minutes after the CPT (run 1 for 10 minutes). The participants then completed the remaining 2 runs of TPI in the MRI scanner for 20 minutes. The fourth saliva sample (S4) and the PANAS (PA4 and NA4) were measured 50 minutes after the CPT (**Fig 1A**).

### Stress Manipulation

Stress induction involved a cold pressor task (CPT) wherein participants submerged their left hand to the wrist in 0–4 °C ice water for 3 consecutive minutes(Riccio et al., 1992). The participants in the control group submerged their left hand in warm water (35–37 °C) for 3 consecutive minutes.

### Third-party Intervention Task

This task included three players. Player A (proposer) received an endowment of 100 MUs per round and could decide how to distribute it between himself/herself and player B (recipient) in units of 5 MUs (i.e., 0, 5, 10, 15, and 20). Player B was required to passively accept the proposal. Player C (the third party) received an endowment of 50 monetary units (MUs, 10 MUs = 1 Chinese yuan) per round and was instructed to observe the collection between player A and player B. In the fMRI scanner, participants believed that they were randomly assigned to player C via a massive drawing process. Participants had three options: transferring MUs to reduce player A’s MUs, transferring MUs to increase player B’s MUs, or keeping all the MUs for themselves. If participants wanted to reduce player A’s money (or increase player B’s money), they needed to determine how many MUs to move from their own 50 MUs and deduct the proposer’s Mus in units of 5 MUs. Following that, the participant could continue to decide how any MUs to move from their remaining MUs to compensate the recipient (**Fig 1E**). The number of trials was preprogrammed to 30 trials for one run, for a total of 3 runs. The offers were created specifically (i.e., The average offer ratio was 90/10 80/20, 70/30, 60/40, and 50/50, but the actual offers shown to the participants fluctuated between 1% and 2%, e.g., 91/9 and 88/12.

### Physiological measures

Saliva cortisol samples were obtained from participants at four time points before and after the stress/control manipulation (Sarstedt, Rom-melsdorf, Germany). These samples were stored at -80 °C until the analysis. We also measured testosterone and oxytocin levels in the saliva to rule out other hormonal effects (Bartz et al., 2011)(see SI in appendix). The samples were thawed and centrifuged for 5 minutes at 3500 rpm. An electrochemiluminescence immunoassay was used to measure salivary cortisol concentrations (Cobas e 601, Roche Diagnostics, Numbrecht, Germany). The intra- and interassay variations for cortisol were below 10%.

Furthermore, as a marker of sympathetic nervous system activity, heart rate was continuously monitored in the CPT stage (3 min) with a Polar WearLink + heart rate monitor (POLAR RCX3) to determine the effects of the stress induction versus the control task; heart rate was also monitored for 3 minutes as a baseline before the CPT (**Fig 1A**).

### Data analysis

#### Behavioral data analysis

To assess the efficacy of the stress manipulation, mixed two-way ANOVAs were performed on salivary cortisol, heart rate, and subjective affective state, with stress manipulation (stress vs. control) as the between-subjects variable and “time” point of measurement as the within-subject factor.

In the third-party intervention task, the rates and costs that subjects chose to punish A, help B or serve themselves were determined in five fair conditions. Since selfish choices were made only in the 50:50 fair condition (**Figure S1**), we focused on the trade-off between help and punishment choices in the subsequent analysis. The number of MUs and rates of punishing A or helping B were subjected to mixed three-way repeated-measures ANOVA with stress manipulation as a between-subjects factor and option (help, punish) and fair condition (50:50, 60:40, 70:30, 80:20, 90:10) as two within-subject factors. Furthermore, we combined 6:4 and 7:3 as relatively unfair conditions (RUF), 80:20 and 90:10 as unfair conditions (UF), and 50:50 as the fair condition (F). The clustering criterion was established based on the findings of a previous meta-analysis(Oosterbeek et al., 2004). Repeated measurement ANOVA was used in two conditions (RUF and UF).

The Greenhouse‒Geisser correction was applied when the requirement of sphericity in the ANOVA for repeated measures was violated. Effect sizes are expressed as partial eta-squared (η ^2^). Behavioral data were analyzed using MATLAB R2019a (Mathworks) and SPSS 25 (IBM).

#### Computational modeling procedures

To better understand the neural correlates underlying acute stress on third-party intervention behaviors, we fitted behavioral data to computational modeling, allowing us to assess the personal involvement of inequality aversion from the perspective of unrelated third parties. We separated the subcomponents of the extent of help (i.e., how averse an individual is to observing someone else being hurt) and punishment severity (i.e., how averse an individual is to observing someone else hurt others), denoted by parameters *β* and *α,* respectively. The values of the two parameters ranged from 0 to 1, with values close to 1 indicating greater magnitude of punishment or help. This model was referred to as the “Other-regarding inequality aversion model” since it hypothesizes that participants’ actions were influenced by aversion to the inequity between the violator and the recipient. The model was formalized as follows:

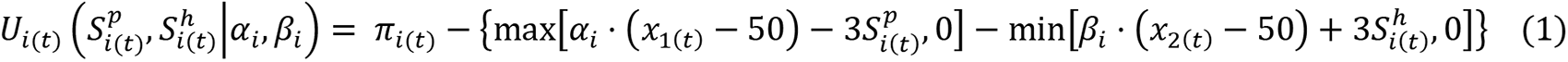

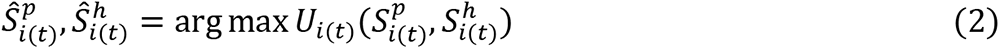

where *i* indicates individual; *U_i_* is Participant *i*’s decision utility of spending tokens on punishment and help at trial *t*; *S^p^*represents the amount of tokens participant used to punish the violator (*x_1_*) and *S^h^* represents the amount of tokens participant used to help the victim (*x_2_*); *π* represents participant’s payoff, in each trial *π* = 50 – S^p^ – S^h^. In Equation 2, 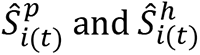 is the number of tokens that optimizes the participant’s subjective utility. Note that 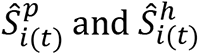 depends not only on the observed level of punishment and help but also on the participant-specific parameters *α_i_* and *β_i_*, which are estimated by minimizing the sum of squared errors (RSS):

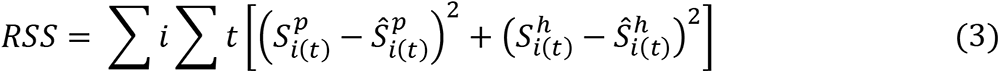

#### Model comparisons

To assess whether the other-regarding inequality aversion model provided a compelling explanation of participants’ behavior, here, three alternative models were compared by calculating the AIC (citation) for each model for each participant.

##### Model 1: Baseline model

The baseline model proposes a rational assumption that participants made decisions only by maximizing their self-payoff. That is, the absence of inequality aversion hypotheses and participant choice not to help and punish instead would be an optimized option. The utility function was as follows:

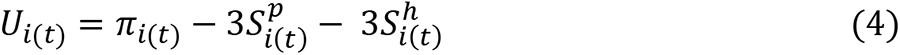

##### Model 2: Self-regarding inequality aversion model

This model assumed that participants made decisions by weighting the payoff inequality between themselves and others versus the violator (who always has more than or equal to 50 tokens), with participants confronting disadvantageous inequality aversion while comparing with receiver participants facing advantageous inequality aversion. The participants’ goal was to eliminate inequality aversion by both punishment and help. The utility function was formalized as follows:

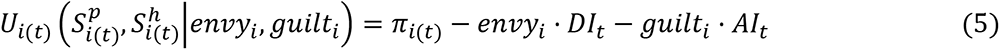

where DI is disadvantageous inequality and AI is advantageous inequality aversion, which is calculated as follows:

Note that in each trial π = 50 – Sp – Sh.

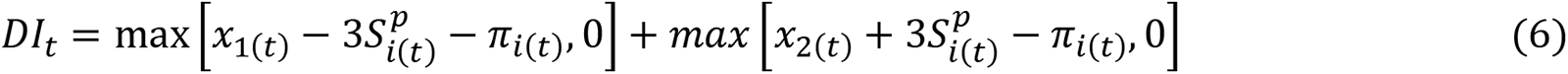

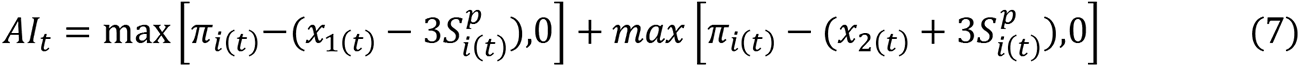

##### Model 3: Other-regarding inequality aversion model with shared parameters for estimating the severity of punishment and help

This model is similar to the other-regarding inequality aversion model introduced above, except that a single parameter denoted the magnitude of punishment and help, *α*, which participants hold the same level of “how averse an individual is to observe someone else being hurt” and “how averse an individual is to observing someone else hurt others”. In other words, participants used the same but distinct psychological mechanisms when determining whether to help or punish others.

#### Model validation

##### Parameter recovery

We ran parameter recovery analyses to ensure that our model was robustly identifiable. This approach allowed us to assess whether our model overfitted the data and to estimate the reliability of the model parameters. To this end, we created 52 simulated participants by simulating task data at random points within the parameter space and fit our model to these simulated subjects’ behaviors.

##### Model prediction

We calculated the correlation between behavioral output predicted by our model and true behaviors. To this end, we first used the estimated parameters of 52 participants to generate predicted behaviors individually and then calculated the correlation between the number of tokens from actual observation in the task and that from predicted behaviors in both punish and help conditions.

### fMRI data analysis

#### Imaging data acquisition and preprocessing

Brain imaging data were acquired on a 3T Prisma MR scanner (Siemens, Germany). During the tasks, blood oxygen level-dependent (BOLD) signals were acquired with a prototype simultaneous multislice echo-planar imaging (EPI) sequence (echo time, 30 ms; repetition time, 2000 ms; field of view, 224 mm × 224 mm; matrix, 112 × 112; inplane resolution, 2 mm × 2 mm; flip angle, 90 degrees; slice thickness, 2.0 mm; 62 slices; slice orientation, transversal). Field map images were acquired using a vendor-provided Siemens gradient echo sequence (gre field mapping: echo time 1, 4.92 ms; echo time 2, 7.38 ms; repetition time, 620 ms; flip angle, 60 degrees; bandwidth, 565 Hz/pixel) with the same geometry and orientation as the EPI image. A high-resolution 3D T1 structural image (3D magnetization-prepared rapid acquisition gradient echo; 0.5 mm × 0.5 mm × 1 mm resolution) was also acquired. Image preprocessing was conducted using the Statistical Parametric Mapping package (SPM12, RRID: SCR_007037; Welcome Department of Imaging Neuroscience, London, United Kingdom). EPI volumes were realigned to the first volume, corrected for geometric distortions using the field map, coregistered to the T1 image, normalized to a standard template (Montreal Neurological Institute, MNI), resampled to 2 × 2 × 2 mm^3^ voxel size, and spatially smoothed with an isotropic 8 mm full-width at half-maximum Gaussian kernel.

#### General linear model

We regressed the fMRI time series into three general linear models (GLMs) to investigate how acute stress affected the brain’s decision circuitry. We looked for neural activity associated with inequity in the decision stage, i.e., the degree of inequity between the proposer and recipient (GLM_ Inequity) with the first GLM. We also checked the stress manipulation effect. In the second GLM (GLM_ Choice), we aimed to recognize brain regions with behavior associated with punishment and help choices under the unfair condition. The third GLM (GLM_ Utility) sought to identify brain regions coding utility and severity preference between punishment and help during the transfer stage. For the main contrasts, the individual voxel threshold was set to *P* < 0.001. We performed whole-brain corrections for multiple comparisons at the cluster level (*P* _FWE_ < 0.05). Furthermore, since we had an a priori hypothesis that the amygdala was related to the “inequity”, we used small-volume correction (SVC) based on anatomically defined bilateral amygdala regions of interest (ROIs) and corrected *P* _FWE_ <0.05. The anatomical ROIs of the bilateral amygdala were created using the SPM Wake Forest University (WFU) Pickatlas toolbox (www.ansir.wfubmc.edu, version 3.0)(Tzourio-Mazoyer et al., 2002).

In the first model (GLM_ Inequity), the “Inequity” δ was defined as “δ = |MU _Proposer_ – MU _Recipient_|”(Zhong et al., 2016). Four regressors were included in the GLM in the following order: (i) onset of the decision stage at the beginning of each trial when participants saw the distributions, (ii) parametric modulation of the trialwise “Inequity”, (iii) onset of the first transfer stage, and (iv) onset of the second transfer stage. We modeled BOLD responses at these onsets as stick functions. All regressors and six head movement regressors of no interest were convolved with a canonical hemodynamic response function. For each event, onset regressor parameter estimates were obtained, and contrast images of each of the parameters against zero were generated. The obtained images were transferred to a second-level random-effects analysis using a two-sample t test and conjunction analysis to compare the stress and control groups.

In the second model (GLM_ Choice), we focused on the decision window when participants responded to the trials of unfair distribution (8:2 & 9:1). We defined the following two onset regressors of interest: (i) onset of the punishment choice of unfair trials and (ii) onset of the help choice of unfair trials. We also defined the following uninterested regressors: (iii) onset of the punishment transfer of unfair trials, (iv) onset of the help transfer of unfair trials, (v) onset of all the choices of relatively unfair trials (7:3 & 6:4, for punishment, help and keep choice), (vi) all the transfers of relatively unfair trials (7:3 & 6:4, for punishment, help and keep choice), (vii) onset of all the choices of fair trials (5:5, for punishment, help and keep choice), and (viii) onset of all the transfers of fair trials (5:5, for punishment, help and keep choice). The GLM additionally included six movement regressors of no interest, three for translational movements (x, y, z) and three for rotation movements (pitch, roll, yaw). All regressors were convolved with the canonical hemodynamic response function. Individual contrast images (for “Punishment”, “Help”, “Punishment-Help”) were transferred to a second-level random-effects analysis using two-sample t tests to compare the stress and control groups.

In the third model (GLM_ Utility), we computed a GLM with a parametric design to identify brain regions coding “Utility” on the transfer window when participants spent chips (MU) to punish the proposer or help the recipient. Utility was defined in our “**Other-regarding inequality aversion model**”. Four regressors were included in the GLM in the following order: (i) onset of the first transfer stage, (ii) parametric modulation of the trialwise “Utility”, (iii) onset of the decision stage at the beginning of each trial, and (iv) onset of the second transfer stage. Similar to the first model, we modeled BOLD responses at these onsets as stick functions. All regressors and six head movement regressors of no interest were convolved with a canonical hemodynamic response function. The obtained images were transferred to a second-level random-effects analysis using two-sample t tests and conjunction analysis to compare the stress and control groups.

#### gPPI analysis

To investigate whether the functional connectivity of the amygdala differed during help and punishing decisions and whether it was affected by stress, we performed a whole-brain gPPI analysis with the right amygdala as the seed region(Stallen et al., 2018). The location of the right amygdala seed ROI was based on a 6 mm radius sphere centered at the peak activation within the contrasts of stress vs. control (GLM_ Inequity). We estimated a GLM with the following regressors: (i) a physiological regressor (i.e., the entire time series of the seed region over the whole experiment), (ii) a psychological regressor for the onset of the punishment choices, (iii) the PPI regressor for the punishment choices, (iv) a psychological regressor for the onset of the help choices, and (v) a PPI regressor for the help choices. The onset and PPI regressors were convolved with a canonical form of the hemodynamic response. The model also included the six motion parameters as regressors of no interest. Individual contrast images for functional connectivity (“punishment”, “help”, “punishment vs. help”) were transferred to a second-level random-effects analysis using a two-sample t test and one-sample t test.

#### Mediation/moderation analysis

We used mediation analysis to test the hypothesis that stress affects punishment rates by amygdala-vmpfc functional coupling. We constructed a mediation model (Model 5) that specified stress manipulation (stress or control) as an independent variable, amygdala-vmpfc functional connectivity as a mediator, and punishment rate as a dependent variable. In addition, a moderated mediation analysis (Model 8) was conducted to test the hypothesis that punishment bias (*α-β*) could account for the indirect associations of value computation in the activity of the rACC, rPCC and rTPJ with punishment severity bias (punishment contribution minus help contribution), and acute stress moderated the correlation between value computation of the rACC, rPCC and rTPJ and punishment bias. These analyses were performed by the SPSS ‘‘PROCESS,’’ made available by Andrew Hayes (Hayes, 2017). A nonparametric resampling procedure (bootstrapping) with 5,000 samples was used to estimate the significance of the indirect effect. Through bootstrapping, we calculated point estimates of the indirect effects and constructed a 95% confidence interval around each point estimate.

## Author contributions

Conceptualization: Huagen Wang, Chao Liu

Methodology: Huagen Wang, Xiaoyan Wu, Jiahua Xu

Investigation: Ruida Zhu, Xiaoqin Mai, Sihui Zhang, Zhenhua Xu

Visualization: Huagen Wang

Supervision: Shaozheng Qin, Chao Liu, Xiaoqin Mai

Writing—original draft: Huagen Wang

Writing—review & editing: Huagen Wang, Shaozheng Qin, Chao Liu

## Data and materials availability

The dataset for this specific manuscript is available from the corresponding author upon request.

## Supplementary Materials

### Supplementary Text

#### Participants

Fifty-three right-handed **male** volunteers were recruited and randomly assigned to either the stress condition (CPT) or the control condition; Data of one participant was removed from analyses, as he failed to understand the task. The final sample included 52 participants (stress: n = 27, mean age = 23.27 years, *SD* = 2.85 years; control: n = 25, mean age = 23.68 years, *SD* = 2.48 years). (1). In a structured phone interview prior to the experiment, participants who met the following conditions were excluded included hormonal contraception, prescription or drug consumption, smoking, alcohol abuse, a history of chronic disease or mental condition, and a major evaluation within two weeks. Furthermore, participants were instructed to abstain from physical activity, meals, and caffeine consumption for 2 hours prior to the study. (2). All study procedures were approved by the Institutional Review Board of the State Key Laboratory of Cognitive Neuroscience and Learning at Beijing Normal University. All participants signed written informed consent.

#### Experimental procedures

Testing sessions took place in the afternoon (1:30–5:30) to control variability in diurnal cortisol secretion. Upon arrival at the lab, participants were guided to the testing room, where they were given a rundown of the experiment and asked to complete questionnaires (psychological and personality tests, see SI) for 20 minutes. Subsequently, the baseline heart rate (HR1) was recorded for 3 minutes, the saliva sample (S1) and the Positive and Negative Affect Scale (PANAS, PA1, and NA1) were measured. Then they were randomly assigned to CPT or control conditions. Heart rate (HR2) was recorded across the whole CPT for 3 minutes. After the CPT or a control manipulation, the saliva sample (S2) and the PANAS (PA2 and NA2) were collected immediately. Then, after 10 minutes’ rest and 10 minutes’ T1 structural image collection in the MRI scanner, the subjects completed the first run of the Third-Party Intervention task (TPI). The third saliva sample (S3) and the PANAS (PA3 and NA3) were measured 25–30 minutes after the CPT (run 1 for 10 minutes). And the participants finished their remaining 2 runs of TPI in the MRI scanner for 20 minutes. The fourth saliva sample (S4) and the PANAS (PA4 and NA4) were measured 50 minutes after the CPT.

#### Stress Manipulation

After the baseline saliva sample was collected, participants were randomly assigned to either a stress or control condition. Stress induction involved a cold pressor task (CPT) wherein participants submerged their left hand to the wrist in 0–4°C ice water for 3 consecutive minutes^1^. If participants failed to complete the CPT, they were excluded from the study. The participants in the control group submerged their left hand in warm water (35–37°C) for 3 consecutive minutes. The CPT has been shown to reliably activate the sympathetic nervous system (SNS) and the hypothalamic-pituitary-adrenal (HPA) axis, as demonstrated by elevations in physiological and endocrine (i.e., cortisol) responses, and it has been used to evoke a stress response.

#### Saliva Cortisol Collection, Storage and Analysis

Saliva samples were obtained from participants at four time points before and after the stress/control manipulation to determine stress-induced changes in cortisol concentrations (Sarstedt, Rom-melsdorf, Germany).

The participants were required to hold a saliva-collecting swab in their mouth for approximately 3 minutes until the saliva completely soaked the swab. Saliva samples were held at -80°C until they were analyzed. To rule out other hormonal effects correlated with prosocial activity, we measured testosterone and oxytocin levels in saliva. The samples were thawed and centrifuged for 5 minutes at 3500 rpm. The concentrations of salivary cortisol were analyzed using electrochemiluminescence immunoassay (Cobas e 601, Roche Diagnostics, Nümbrecht, Germany), and those of testosterone were analyzed using an enzyme-linked immunoassay kit developed for saliva (Salimetrics, State College, PA) with sensitivity of 0.500 nmol/L (lower limit). The standard range in the assay was 0.5–1,750 nmol/L for cortisol. The intra- and inter-assay coefficient variations for cortisol were below 10%, and we did not test the precision of testosterone and oxytocin because there was not enough saliva.

#### Heart rate Collection

Furthermore, as markers of sympathetic nervous system activity, heart rate was continuously monitored in the CPT stage (3 min) with a Polar WearLink + heart rate monitor (POLAR RCX3) to determine the effects of the stress induction versus the control task, and heart rate was also monitored for 3 minutes as a baseline before the CPT.

#### Psychological measures

The Positive and Negative Affect Scale (PANAS) was used to measure the subjective affective states of participants at each designated instant. The scale has a total of 20 items describing different feelings and emotions, including 10 items for positive affects (e.g., “interested”, “excited”) and 10 items for negative affect (e.g., “nervous”, “scared”). The participants were asked to score each item on a 5-point scale based on their instant affective state, from 1 (very slightly or not at all) to 5 (extremely). The average scores of positive affect (PA) and negative affect (NA) were calculated. Besides, studies have shown that some personality traits (such as impulsiveness) and empathy^2,3^ also have a very important impact on prosocial behavior, therefore, some personality factors were also considered in this study. Before the experiment, participants completed an online survey (implemented via Qualtrics software, 2009, Provo, UT, USA. https://www.qualtrics.com), which included demographic questions and some personality measures: Perceived Stress Scale (PSS)^4^; State-Trait Inventory for Cognitive and Somatic Anxiety (STICSA)^5^; Social Value Orientation (SVO)^6^; Behavioral Inhibition System and Behavioral Activation System Scales (BIS/BAS Scales)^7^; Interpersonal Reactivity Index (IRI)^8^; Altruistic personality scale (APS)^9^.

#### Third-party Intervention Task

This task includes three players. The player A (proposer) received an endowment of 100 MUs per round and could decide how to distribute it between himself/herself and the player B (recipient) in units of 5-MUs (i.e., 0, 5, 10, 15, and 20).The player B had to passively accept the proposal. The player C (the third party) received an endowment of 50 monetary units (MUs, 10 MUs = 1 Chinese yuan) per round and was instructed to observe the collection between player A and player 0. B. In the fMRI scanner, participants believed that they were randomly assigned to player C via a massive drawing process. Participants had three options: transferring MUs to reduce player A’s MUs, transferring MUs to increase player B’s MUs, or keeping all the MUs for themselves. If participants wanted to reduce player A’s money (or increase player B’s money), they needed to determine how many MUs to move from their own 50 MUs and deduct the proposer’s Mus in units of 5 MUs. Following that, he can continue to decide how much MUs to move from their remaining MUs to compensate the recipient. The number of trials was pre-programmed to 30 trials for one run, a total of 3 runs. The offers were created specifically (i.e. the average offer ratio was 90/10 80/20, 70/30, 60/40, 50/50, but the actual offers shown to the participants fluctuated between 1% and 2%, e.g. 91/9, 88/12).

There are a few specifics to note: 1) When player C decided to deduct A’s MUs or increase B’s MUs, the cost ratio was 1:3, as previously reported^2^, which means that every MU transferred by player C can be deducted or increased by 3 MUs to player A or player B, respectively. 2) In the expression of instructions, we used “player A, B, and C” instead of “dictator”, “recipient” and “observer” and “deduct” and “increase” in place of “punish” and “help” 3) Player A and B were not real; we had pre-programmed the allocation chosen by player A. 4) For each participant, the order of the trials in each run was randomized.

Functional Magnetic Resonance Imaging (fMRI) Procedure

#### Imaging data acquisition and preprocessing

Brain imaging data were acquired on a 3T Prisma MR scanner (Siemens, Erlangen, Germany) with a 64-channel phased-array head-neck coil for signal reception. During the tasks, blood oxygen level-dependent (BOLD) signals were acquired with a prototype simultaneous multi-slice echo-planar imaging (EPI) sequence (echo time, 30 ms; repetition time, 2000 ms; field of view, 224 mm × 224 mm; matrix, 112 × 112; inplane resolution, 2 mm × 2 mm; flip angle, 90 degree; slice thickness, 2.0 mm; gap, 15%; the number of slices, 62; slice orientation, transversal; bandwidth, 2232 Hz/Pixel; slice acceleration factor, 2). Field map images were acquired using a vendor-provided Siemens gradient echo sequence (gre field mapping: echo time 1, 4.92 ms; echo time 2, 7.38 ms; repetition time, 620 ms; flip angle, 60 degree; bandwidth, 565 Hz/Pixel) with the same geometry and orientation as the EPI image. A high-resolution 3D T1 structural image (3D magnetization-prepared rapid acquisition gradient echo; 0.5 mm × 0.5 mm × 1 mm resolution) was also acquired. Image preprocessing was performed using the Statistical Parametric Mapping package (SPM12, RRID: SCR_007037; Welcome Department of Imaging Neuroscience, London, United Kingdom). EPI volumes were realigned to the first volume, corrected for geometric distortions using the field map, coregistered to the T1 image, normalized to a standard template (Montreal Neurological Institute, MNI), resampled to 2 × 2 × 2 mm^3^ voxel size, and spatially smoothed with an isotropic 8 mm full-width at half-maximum Gaussian kernel.

#### fMRI data analysis

During the conversion process, the first three images at the beginning of the functional runs were discarded to enable the signal to achieve steady-state equilibrium between radio frequency pulsing and relaxation. Images were motion corrected for three translational and three rotational directions.

#### General linear model

We regressed the fMRI time series into three general linear models (GLMs) to investigate how acute stress affected the brain’s decision circuitry. We looked for neural activity associated with inequity on the decision stage, i.e., the degree of inequity between the proposer and recipient (GLM_ Inequity) with the first GLM, and in this model, we also check the stress manipulation effect. In the second GLM (GLM_ Choice), we aimed to recognize brain regions whose behavior was associated with punishment and help choice under the unfair condition. The third GLM (GLM_ Utility) sought to identify brain regions coding utility and severity preference between punishment and help during the transfer stage. For the main contrasts, the individual voxel threshold was set to P < 0.001. We performed whole-brain corrections for multiple comparisons at the cluster level (P FWE < 0.05). Furthermore, since we had an a priori hypothesis that the amygdala was related to the “inequity,” we used small volume correction (SVC) based on anatomically defined bilateral amygdala region of interests (ROIs) and corrected P FWE <0.05. The anatomical ROIs of the bilateral amygdala were created using the SPM Wake Forest University (WFU) Pickatlas toolbox (www.ansir.wfubmc.edu, version 3.0).

##### In the first model (GLM_ Inequity)

We computed a GLM with a parametric design to identify brain regions coding “Inequity” on the decision window when participants were informed how many MUs had been distributed by the proposer. In each trial, the “Inequity” δ was defined as “δ = |MU Proposer – MU Recipient|”48. Four regressors were included in the GLM in the following order: (i) onset of the decision stage at the beginning of each trial when participants saw the distributions, (ii) parametric modulation of the trial-wise “Inequity”, (iii) onset of the first transfer stage, and (iv) onset of the second transfer stage. We modeled BOLD responses at these onsets as stick functions. All regressors and six head movement regressors of no interest were convolved with a canonical hemodynamic response function. For each event, onset regressor parameter estimates were obtained and contrast images of each of the parameters against zero were generated. The obtained images were transferred to a second-level random-effects analysis using two-sample t-test and conjunction analysis to compare the stress and control groups.

##### In the second model (GLM_ Choice)

We focused on the decision window when participants responded to the trials of unfair distribution (8:2 & 9:1). We defined the following two onset regressors of interest: (i) onset of the punishment choice of unfair trials, (ii) onset of the help choice of unfair trials. We also defined the following uninterested regressors: (iii) onset of the punishment transfer of unfair trials, (iv) onset of the help transfer of unfair trials, (v) onset of all the choices of relative unfair trials (7:3 & 6:4, for punishment, help and keep choice), (vi) all the transfers of relative unfair trials (7:3 & 6:4, for punishment, help and keep choice), (vii) onset of all the choices of fair trials (5:5, for punishment, help and keep choice), (viii) onset of all the transfers of fair trials (5:5, for punishment, help and keep choice). The GLM additionally included six movement regressors of no interest, three for translational movements (x, y, z) and three for rotation movements (pitch, roll, yaw). All regressors were convolved with the canonical hemodynamic response function. Individual contrast images (for “Punishment”, “Help”, “Punishment-Help”) were transferred to a second-level random-effects analysis using two-sample t-tests to compare the stress and control groups. More specifically, several participants were excluded in the second level because coefficients for the parameters could not be estimated when participants never, or only one time, choosing help or punishment per functional run. To this end, the group contrast of “Punishment-Help” was computed with 43 participants (stress group: 22; and control group:21), the group contrast of “Punishment” was computed with 50 participants (stress group: 26; and control group:24), the group contrast of “Help” was computed with 45 participants (stress group: 23; and control group:22).

##### In the third model (GLM_ Utility)

We computed a GLM with a parametric design to identify brain regions coding “Utility” on the transfer window when participants spending chips (MU) to punish the proposer or help the recipient. The Utility was defined in our “Other-regarding inequality aversion model”. Four regressors were included in the GLM in the following order: (i) onset of the first transfer stage, (ii) parametric modulation of the trial-wise “Utility”, (iii) onset of the decision stage at the beginning of each trial, and (iv) onset of the second transfer stage. Same as the first model, we modeled BOLD responses at these onsets as stick functions.

All regressors and six head movement regressors of no interest were convolved with a canonical hemodynamic response function. For each event, onset regressor parameter estimates were obtained and contrast images of each of the parameters against zero were generated. The obtained images were transferred to a second-level random-effects analysis using two-sample t tests and conjunction analysis to compare the stress and control groups.

#### Functional connectivity analysis (gPPI analysis)

To investigate whether the functional connectivity of the amygdala differed during help and punishing decisions and whether it was affected by stress, we performed a whole-brain gPPI analysis with the right amygdala as seed region5. The location of the right amygdala seed ROI was based on a 6 mm radius sphere centered at the peak activation within the contrasts of stress vs. control (GLM_ Inequity). We estimated a GLM with the following regressors: (i) a physiological regressor (i.e., the entire time series of the seed region over the whole experiment), 0. (ii) a psychological regressor for the onset of the punishment choices, (iii) the PPI regressor for the punishment choices, (iv) a psycho-logical regressor for the onset of the help choices, and (v) a PPI regressor for the help choices. The onset and PPI regressors were convolved with a canonical form of the hemodynamic response. The model also included the six motion parameters as regressors of no interest. Individual contrast images for functional connectivity (“punishment”, “help”, “punishment vs. help”) were transferred to a second-level random-effects analysis using a two-sample t test and one-sample t test. We performed whole-brain corrections for multiple comparisons at the cluster level (P FWE < 0.05), the individual voxel threshold was set to P < • All reported coordinates (x, y, z) are in MNI space.

---

**Fig. S1.**
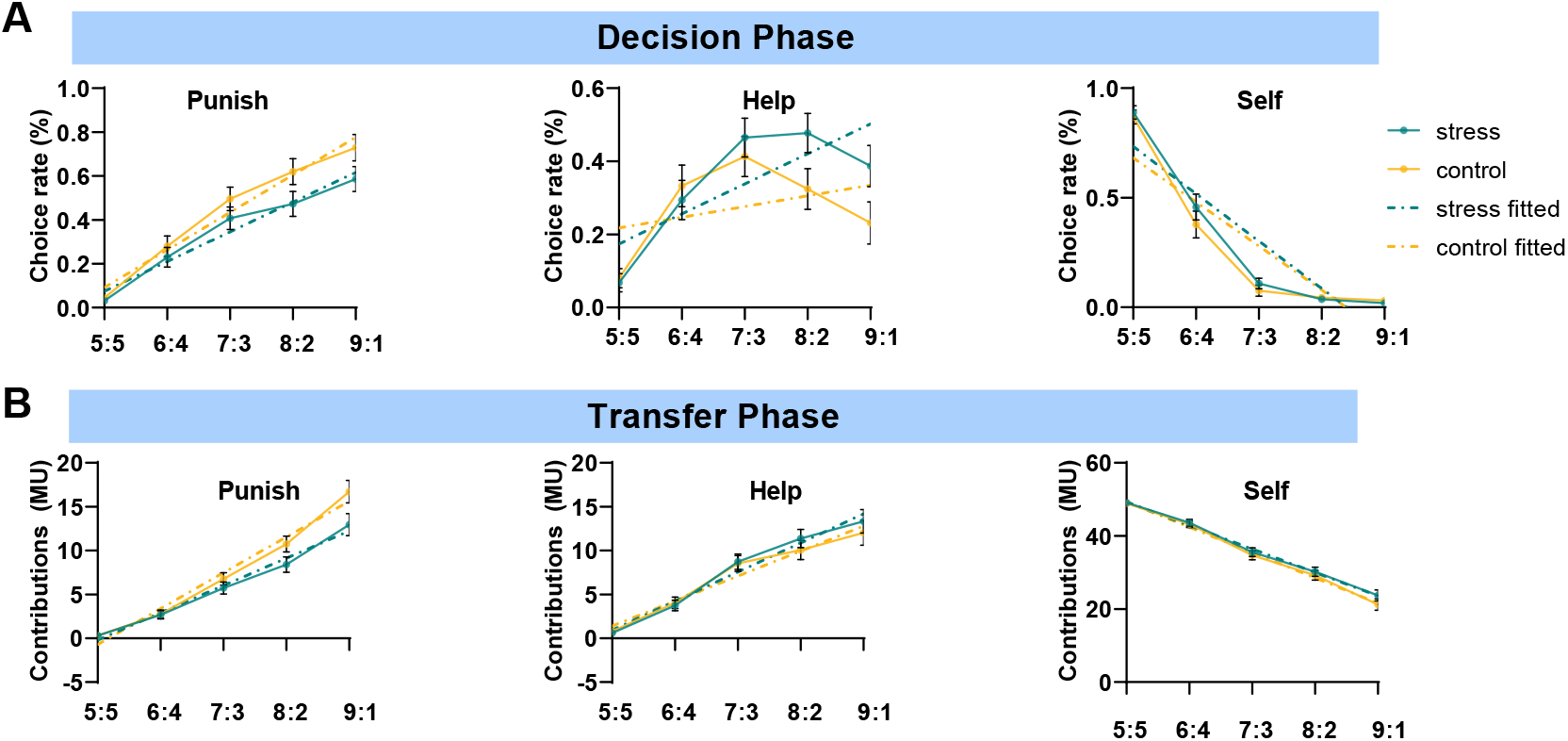
Acute stress modulated the prosocial preference. We compared the punishment, help and selfish choices, as a function of the fair condition (50:50, 60:40, 70:30, 80:20, 90:10). As expected, we found that as the degree of unfairness increased, the proportion of punishment and help behavior increased, and the proportion of selfish behavior decreased correspondingly and this effect was similar in the stress and control group (**Fig S1A&B**, stress main effect of choice rate: F (1, 50) = 0.71, P = 0.40, *η*p^2^ = 0.014; stress main effect of contribution: F (1, 50) = 1.04, P = 0.31, *η*p^2^ = 0.020).

**Fig. S2.**
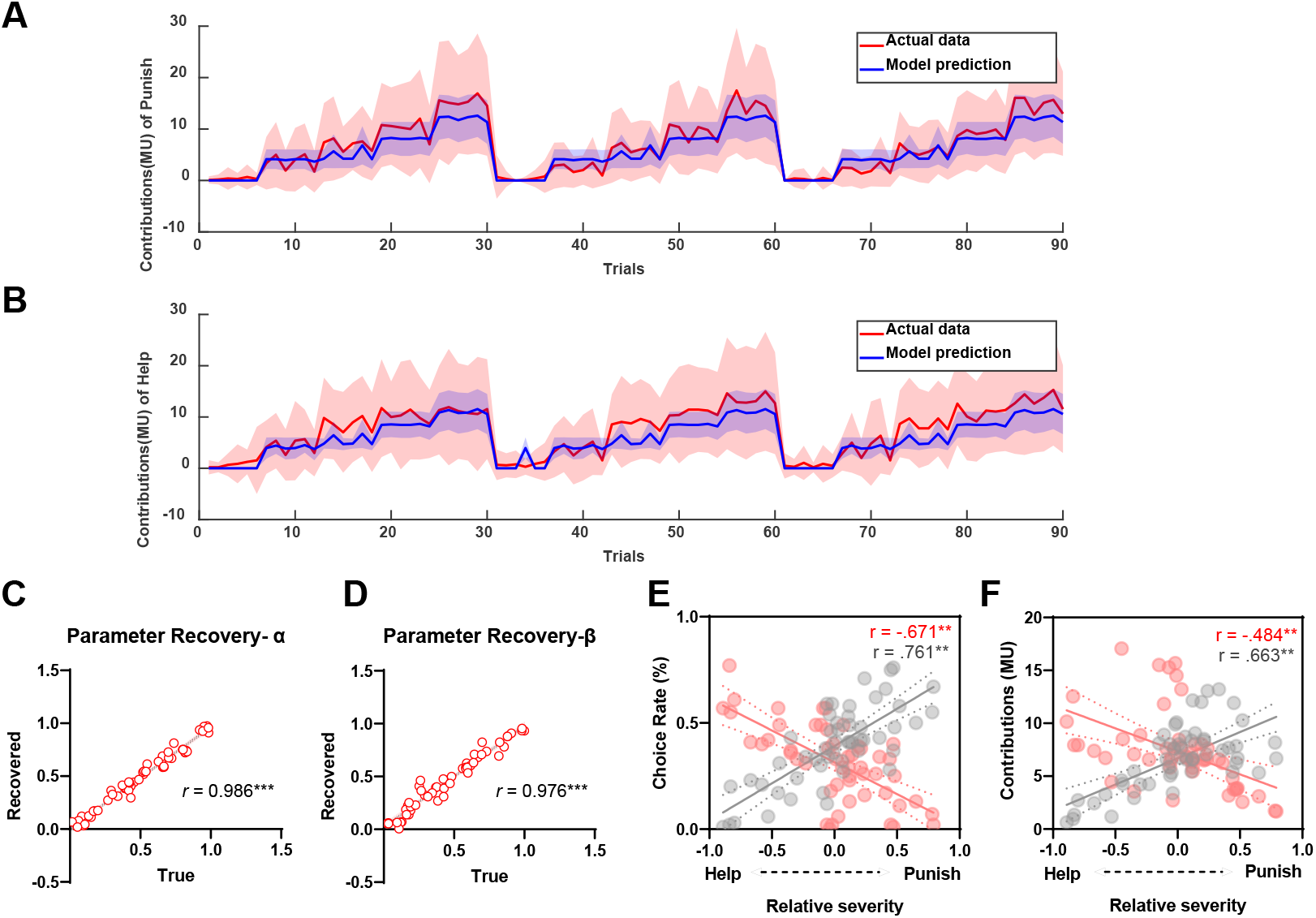
Model validation and Parameter recovery. The posterior predictive check further showed a high correlation between the actual behaviors and model prediction (**Fig S2 A&B**, r = 0.82, p < 0.001 in punishment contribution and r = 0.75, p < 0.001 in help contribution), which suggests that our model can predict participants’ behavior well. The recovery analysis revealed that the correspondence between the true and recovered parameters was significantly high for two free parameters (**Fig S2 C&D,** *α*: *r* = 0.986, *p* < 0.001; *β*: *r* = 0.976, *p* < 0.001). Furthermore, the model predicted punishment bias (difference of the two free parameters: α-β) was positively related to the actual punishment rate and contributions, (**Fig S2E& S2F**, r = 0.76, *P* < 0.01 for punishment rate and r = 0.66, *P* < 0.01 for punishment contributions), but negatively related to the actual help rate and contributions (r = -0.67, *P* < 0.01 for help rate and r = -0.48, *P* < 0.01 for help contributions).

**Fig. S3.**
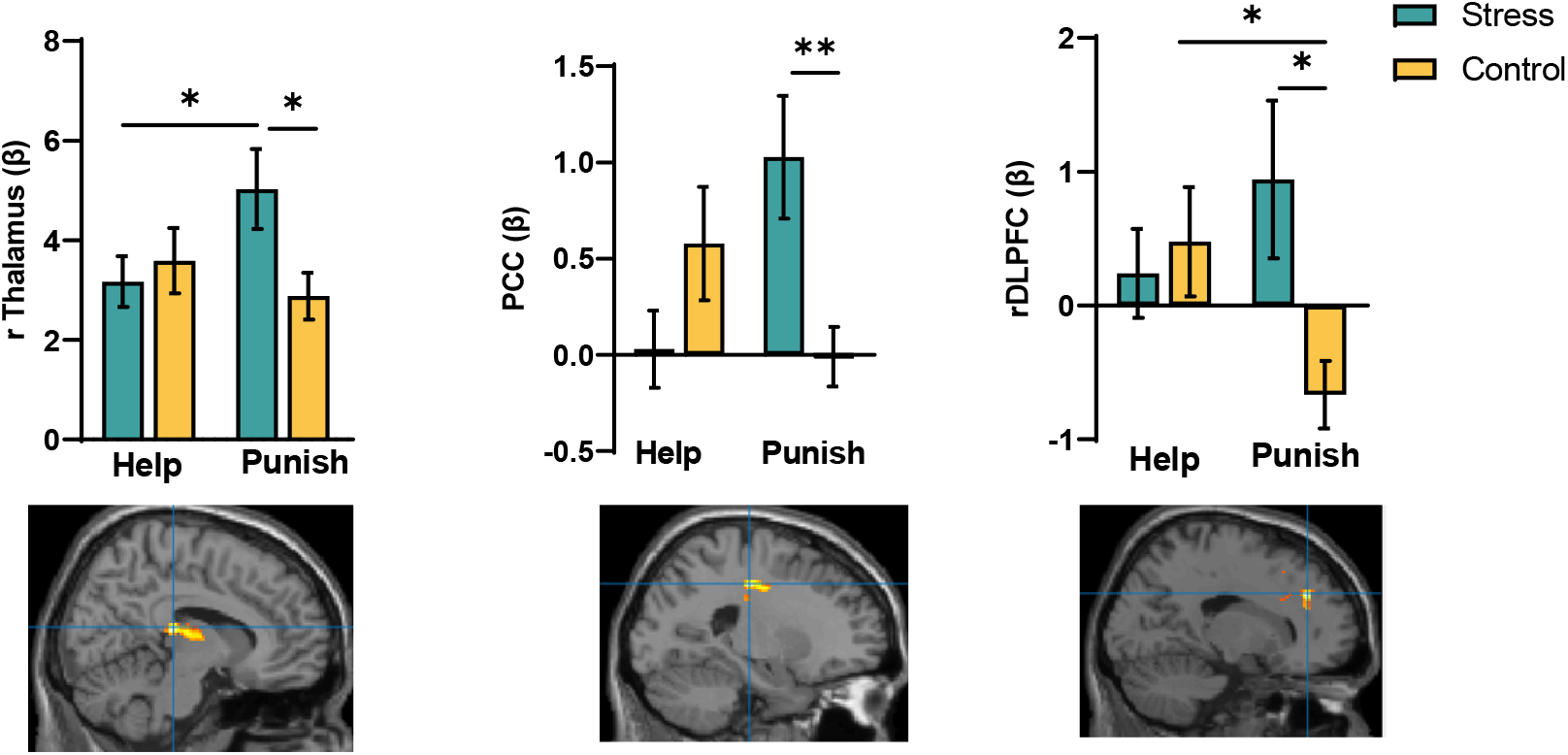
Stress influence the neural correlates of punishment versus help decision under all unfair conditions (relative and unfair condition, 60:40∼90:10). We also regressed the fMRI time series into a general linear model (GLM_ Choice2) to investigate how acute stress affected the brain’s decision circuitry. In the GLM (GLM_ Choice2), we aimed to recognize brain regions whose behavior was associated with punishment and help choice under all the unfair conditions (60:40∼90:10). We defined the following two onset regressors of interest: (i) onset of the punishment choice of all unfair trials, (ii) onset of the help choice of all unfair trials. We also defined the following uninterested regressors: (iii) onset of the punishment transfer of all unfair trials, (iv) onset of the help transfer of all unfair trials, (v) onset of all the choices of fair trials (5:5, for punishment, help and keep choice), (vi) onset of all the transfers of fair trials (5:5, for punishment, help and keep choice). The GLM additionally included six movement regressors of no interest, three for translational movements (x, y, z) and three for rotation movements (pitch, roll, yaw). All regressors were convolved with the canonical hemodynamic response function. Individual contrast images (for “Punishment”, “Help”) were transferred to a second-level analysis using 2 (group) by 2 (Choice) mixed factorial analysis of variance (ANOVA)^11^. More specifically, several participants were excluded in the second level because coefficients for the parameters could not be estimated when participants never, or only one time, choosing help or punishment per functional run. To this end, the group contrast was computed with 48 participants (stress group: 25; and control group:23), By comparing trials in which punishment is selected vs trials in which help is selected between stress and control group under all unfair conditions, we found that stress induced a higher activity in right DLPFC, PCC and right Thalamus when select punishment choices relative to help choices (Figure S3, initial threshold set at *P* uncorrected < 0.001, whole-brain cluster corrected at P FWE < 0.05).

**Table S1.** Control variables per group [M (SD)].

|  | Stress | Control |
| --- | --- | --- |
| PSS | 25.38(1.43) | 24.12(0.99) |
| STICSA | 16.40(0.97) | 15.66(0.96) |
| SVO | 34.64(1.63) | 34.92(1.66) |
| BIS | 2.65(0.10) | 2.73(0.07) |
| BAS | 2.94(0.10) | 3.13(0.08) |
| IRI | 2.52(0.07) | 2.58(0.06) |
| APS | 58.35(3.54) | 63.64(2.92) |
| Base Testosterone | 2.06(0.14) | 2.16(0.12) |
| Base Oxytocin | 17.20(0.83) | 17.69(0.84) |

**Psychological and personality measures:** Before the experiment, participants completed an online survey (implemented via Qualtrics software, 2009, Provo, UT, USA. https://www.qualtrics.com), which included demographic questions and some personality measures: Perceived Stress Scale (PSS)<sup>4</sup>; State-Trait Inventory for Cognitive and Somatic Anxiety (STICSA)<sup>5</sup>; Social Value Orientation (SVO)<sup>6</sup>; Behavioral Inhibition System and Behavioral Activation System Scales (BIS/BAS Scales)<sup>7</sup>; Interpersonal Reactivity Index (IRI)<sup>8</sup>; Altruistic personality scale (APS)<sup>9</sup>. We found no group differences in any of these variables (all $P > 0.131$ ).

**Table S2.**
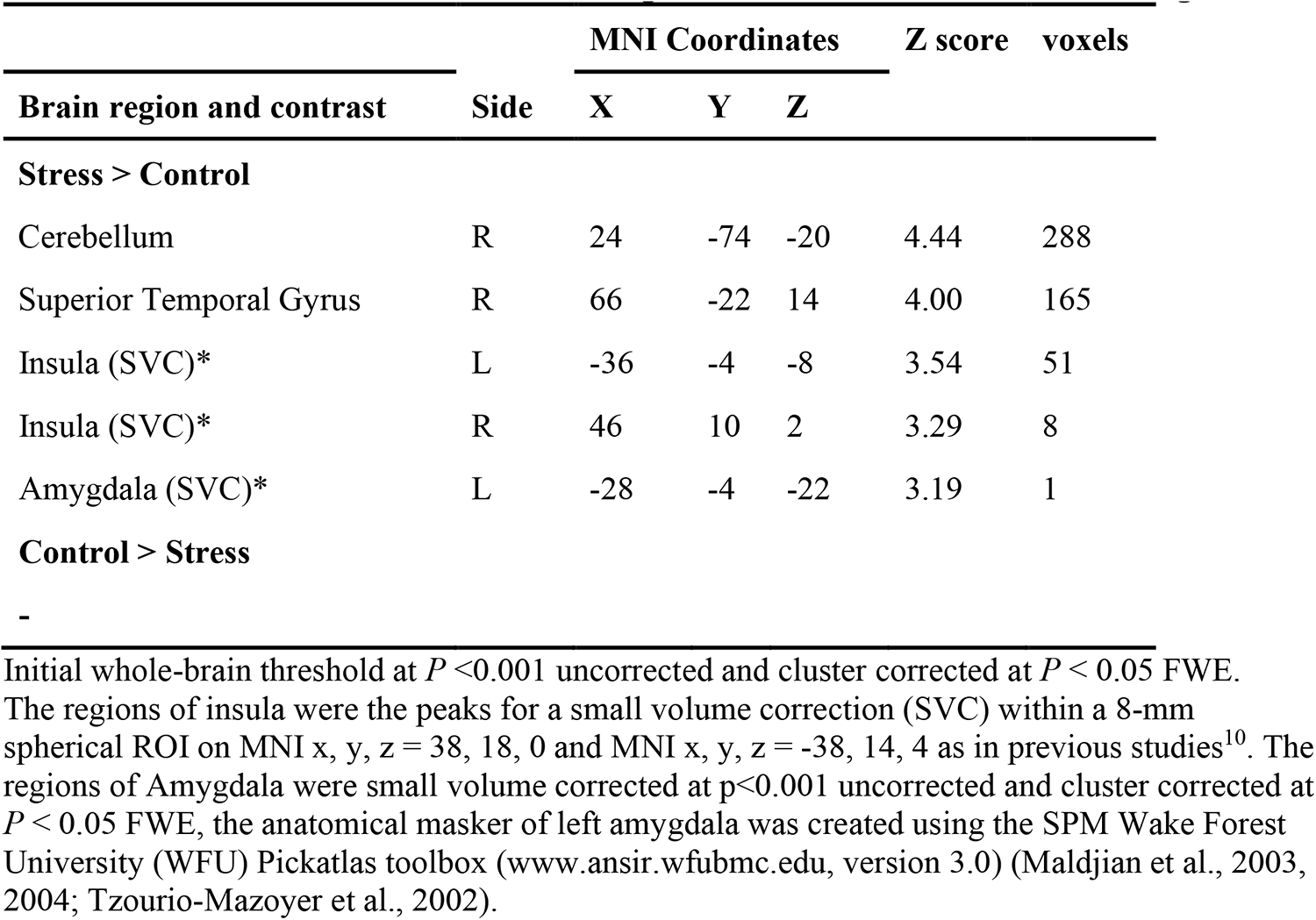
Stress-induced differences in the brain among all fair conditions in the decision stage.

| Brain region and contrast | Side | MNI Coordinates |  |  | Z score | voxels |
| --- | --- | --- | --- | --- | --- | --- |
|  |  | X | Y | Z |  |  |
| **Stress > Control** |  |  |  |  |  |  |
| Cerebellum | R | 24 | -74 | -20 | 4.44 | 288 |
| Superior Temporal Gyrus | R | 66 | -22 | 14 | 4.00 | 165 |
| Insula (SVC)* | L | -36 | -4 | -8 | 3.54 | 51 |
| Insula (SVC)* | R | 46 | 10 | 2 | 3.29 | 8 |
| Amygdala (SVC)* | L | -28 | -4 | -22 | 3.19 | 1 |
| **Control > Stress** |  |  |  |  |  |  |
| - |  |  |  |  |  |  |

**Table S3.**
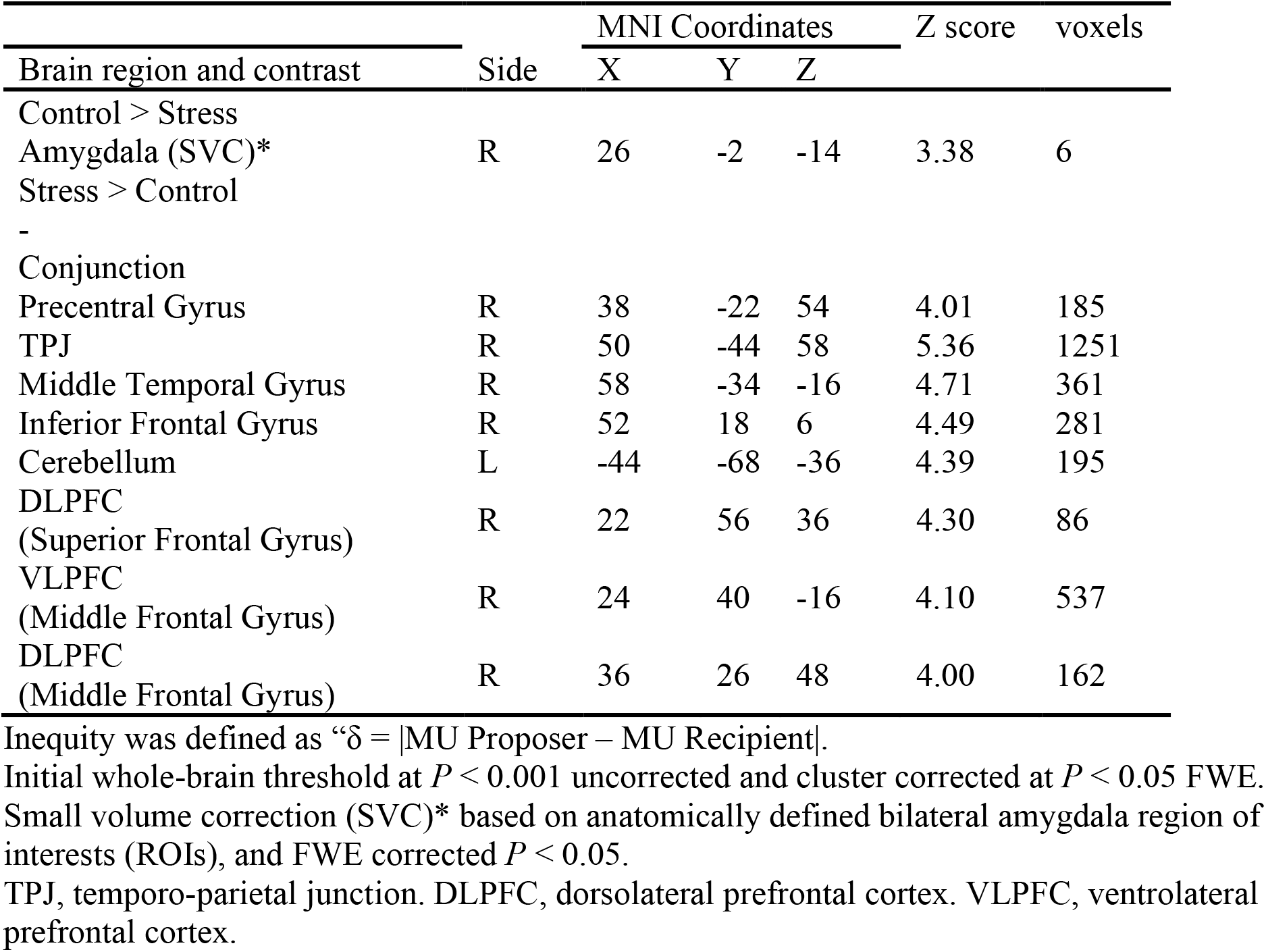
Neural response for increasing distributional inequity |Proposer-Recipient | in decision stage.

**Table S4.** Neural Correlates of utility in Transfer stage.

| Brain region and contrast | Side | MNI Coordinates |  |  | Z score | voxels |
| --- | --- | --- | --- | --- | --- | --- |
|  |  | X | Y | Z |  |  |
| Control > Stress |  |  |  |  |  |  |
| Stress > Control | - |  |  |  |  |  |
| Conjunction |  |  |  |  |  |  |
| Precentral Gyrus | R | 36 | -24 | 54 | 5.25 | 831 |
| Fusiform Gyrus | L | -8 | -66 | -4 | 5.13 | 340 |
| VMPFC<br>(Medial Frontal Gyrus) | L | -6 | 32 | -8 | 4.58 | 1039 |
| Middle Temporal Gyrus | R | 60 | -54 | 12 | 4.40 | 157 |
| Insula | R | 46 | -20 | 20 | 4.38 | 185 |
| Middle Temporal Gyrus | L | -56 | -16 | -8 | 4.36 | 408 |
| Posterior Cingulate | L | -6 | -54 | 18 | 4.34 | 297 |
| Cuneus | R | 12 | -88 | 28 | 4.31 | 654 |
| Superior Temporal Gyrus | L | -60 | -60 | 24 | 4.24 | 312 |
| Middle Temporal Gyrus | R | 64 | -18 | -6 | 3.69 | 180 |
| DLPFC<br>(Middle Frontal Gyrus) | R | 34 | 32 | 28 | 4.30 | 358 |

Initial whole-brain threshold at $P < 0.001$ uncorrected and cluster corrected at $P < 0.05$ FWE.
VMPFC, ventromedial prefrontal cortex. DLPFC, dorsolateral prefrontal cortex.

**Table S5.**
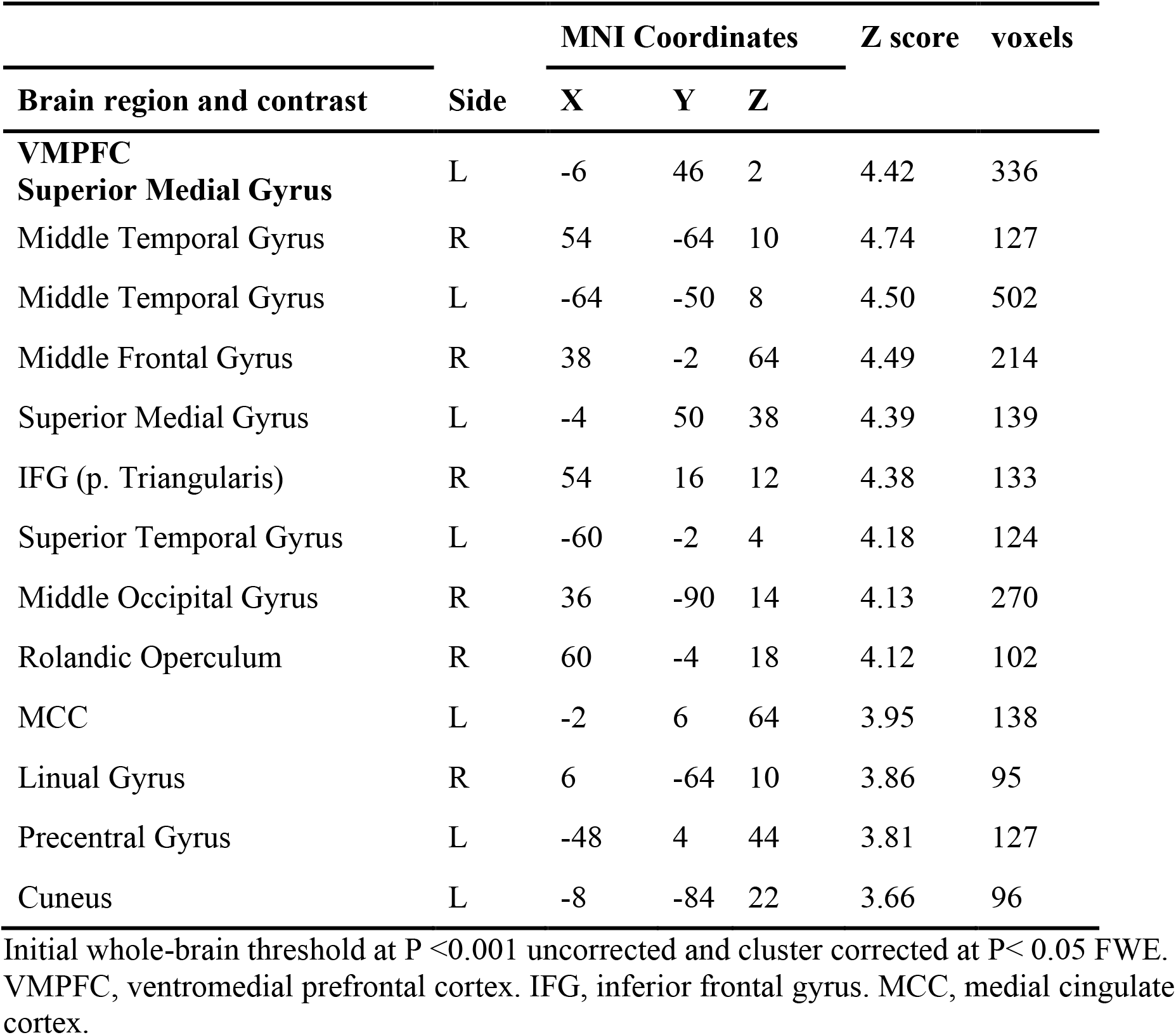
PPI: Regions showing stress group functional connectivity with the right amygdala when making a punishment choice in the unfair condition.

**Table S6.**
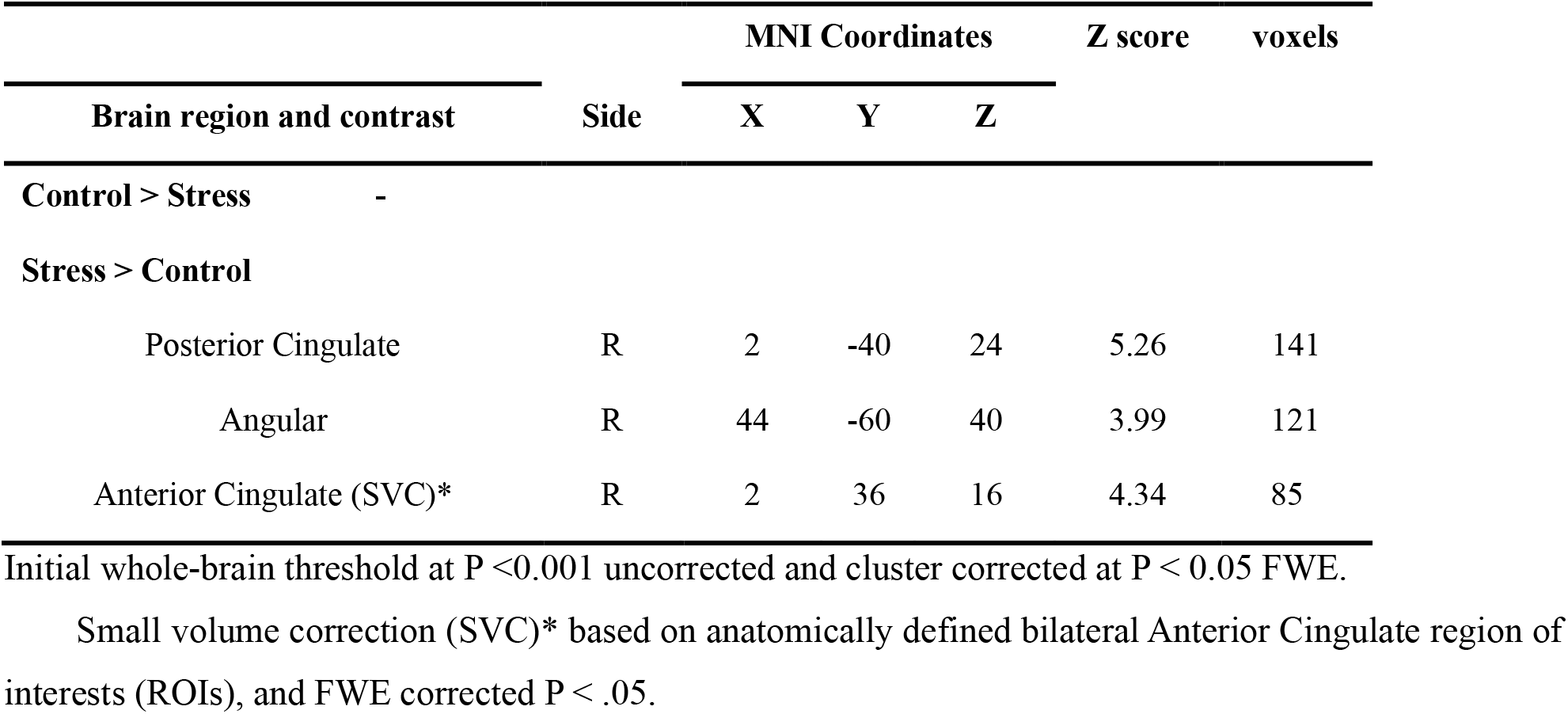
Stress induced higher neural correlates of relative severity in punishment versus help (α-β).

## References

1. Fehr, E., Fischbacher, U. & Gächter, S. Strong reciprocity, human cooperation, and the enforcement of social norms. Human Nature 13, 1–25 (2002).

2. Fehr, E. & Fischbacher, U. Social norms and human cooperation. Trends Cogn Sci 8, 185–190 (2004).

3. Fehr, E. & Fischbacher, U. The nature of human altruism. Nature 425, 785–791 (2003).

4. FeldmanHall, O., Sokol-Hessner, P., Van Bavel, J. J. & Phelps, E. A. Fairness violations elicit greater punishment on behalf of another than for oneself. Nat Commun 5, 1–6 (2014).

5. Stallen, M. et al. Neurobiological Mechanisms of Responding to Injustice. The Journal of Neuroscience 1242–17 (2018).

6. David, B., Hu, Y., Krüger, F. & Weber, B. Other-regarding attention focus modulates third-party altruistic choice: An fMRI study. Sci Rep 7, 43024 (2017).

7. Phelps, E. A., Lempert, K. M. & Sokol-Hessner, P. Emotion and Decision Making: Multiple Modulatory Neural Circuits. Annu Rev Neurosci 37, 263–287 (2014).

8. Ulrich-Lai, Y. M. & Herman, J. P. Neural regulation of endocrine and autonomic stress responses. Nat Rev Neurosci 10, 397–409 (2009).

9. Kahneman, D. Thinking, Fast and Slow. (2013).

10. Hallsson, B. G., Siebner, H. R. & Hulme, O. J. Fairness, fast and slow: A review of dual process models of fairness. Neuroscience and Biobehavioral Reviews vol. 89 49–60 (2018).

11. Hermans, E. J. et al. Dynamic adaptation of large-scale brain networks in response to acute stressors. Trends Neurosci 37, 304–314 (2014).

12. Arnsten, A. F. T. Stress weakens prefrontal networks: molecular insults to higher cognition. Nat Neurosci 18, 1376–1385 (2015).

13. Taylor, S. E. Tend and befriend: Biobehavioral bases of affiliation under stress. Curr Dir Psychol Sci 15, 273–277 (2006).

14. Tomova, L. et al. Increased neural responses to empathy for pain might explain how acute stress increases prosociality. Soc Cogn Affect Neurosci 12, 401–408 (2017).

15. FeldmanHall, O., Raio, C. M., Kubota, J. T., Seiler, M. G. & Phelps, E. A. The Effects of Social Context and Acute Stress on Decision Making Under Uncertainty. Psychol Sci 26, 1918–1926 (2015).

16. Xie, E., Liu, M., Liu, J., Gao, X. & Li, X. Neural mechanisms of the mood effects on third-party responses to injustice after unfair experiences. Hum Brain Mapp (2022) doi:10.1002/hbm.25874.

17. Civai, C., Huijsmans, I. & Sanfey, A. G. Neurocognitive mechanisms of reactions to second- and third-party justice violations. Sci Rep 9, 1–11 (2019).

18. Takahashi, T., Ikeda, K. & Hasegawa, T. Social evaluation-induced amylase elevation and economic decision-making in the dictator game in humans. Neuroendocrinology Letters 28, 662–665 (2007).

19. von Dawans, B., Fischbacher, U., Kirschbaum, C., Fehr, E. & Heinrichs, M. The Social Dimension of Stress Reactivity. Psychol Sci 23, 651–660 (2012).

20. Nickels, N., Kubicki, K. & Maestripieri, D. Sex Differences in the Effects of Psychosocial Stress on Cooperative and Prosocial Behavior: Evidence for ‘Flight or Fight’ in Males and ‘Tend and Befriend’ in Females. Adaptive Human Behavior and Physiology 3, 171–183 (2017).

21. Hu, Y., Strang, S. & Weber, B. Helping or punishing strangers: Neural correlates of altruistic decisions as third-party and of its relation to empathic concern. Front Behav Neurosci 9, 24 (2015).

22. Buchanan, T. W. & Preston, S. D. Stress leads to prosocial action in immediate need situations. Frontiers in Behavioral Neuroscience vol. 8 (2014).

23. Buckholtz, J. W. et al. The Neural Correlates of Third-Party Punishment. Neuron 60, 930–940 (2008).

24. Bartra, O., McGuire, J. T. & Kable, J. W. The valuation system: A coordinate-based meta-analysis of BOLD fMRI experiments examining neural correlates of subjective value. Neuroimage 76, 412–427 (2013).

25. Büchel, C., Holmes, A. P., Rees, G. & Friston, K. J. Characterizing stimulus-response functions using nonlinear regressors in parametric fMRI experiments. Neuroimage (1998).

26. von Dawans, B., Fischbacher, U., Kirschbaum, C., Fehr, E. & Heinrichs, M. The Social Dimension of Stress Reactivity: Acute Stress Increases Prosocial Behavior in Humans. Psychol Sci 23, 651–660 (2012).

27. Nickels, N., Kubicki, K. & Maestripieri, D. Sex Differences in the Effects of Psychosocial Stress on Cooperative and Prosocial Behavior: Evidence for ‘Flight or Fight’ in Males and ‘Tend and Befriend’ in Females. Adaptive Human Behavior and Physiology 3, 171–183 (2017).

28. Youssef, F. F., Bachew, R., Bissessar, S., Crockett, M. J. & Faber, N. S. Sex differences in the effects of acute stress on behavior in the ultimatum game. Psychoneuroendocrinology 96, 126–131 (2018).

29. Dreber, A., Rand, D. G., Fudenberg, D. & Nowak, M. A. Winners don’t punish. Nature (2008) doi:10.1038/nature06723.

30. Rockenbach, B. & Milinski, M. The efficient interaction of indirect reciprocity and costly punishment. Nature 444, 718–723 (2006).

31. Raihani, N. J. & Bshary, R. Third-party punishers are rewarded, but third-party helpers even more so. Evolution (N Y) 69, 993–1003 (2015).

32. Sylwester, K. & Roberts, G. Reputation-based partner choice is an effective alternative to indirect reciprocity in solving social dilemmas. Evolution and Human Behavior 34, 201–206 (2013).

33. Sanfey, A. G., Rilling, J. K., Aronson, J. A., Nystrom, L. E. & Cohen, J. D. The neural basis of economic decision-making in the Ultimatum Game. Science (1979) 300, 1755–1758 (2003).

34. Knoch, D., Pascual-Leone, A., Meyer, K., Treyer, V. & Fehr, E. Diminishing reciprocal fairness by disrupting the right prefrontal cortex. Science (1979) 314, 829–832 (2006).

35. Zinchenko, O. & Klucharev, V. Commentary: The Emerging Neuroscience of Third-Party Punishment. Frontiers in Human Neuroscience vol. 11 499–501 (2017).

36. Pearson, J. M., Heilbronner, S. R., Barack, D. L., Hayden, B. Y. & Platt, M. L. Posterior cingulate cortex: Adapting behavior to a changing world. Trends in Cognitive Sciences vol. 15 (2011).

37. Grueschow, M., Polania, R., Hare, T. A. & Ruff, C. C. Automatic versus Choice-Dependent Value Representations in the Human Brain. Neuron 85, (2015).

38. Schurz, M., Radua, J., Aichhorn, M., Richlan, F. & Perner, J. Fractionating theory of mind: A meta-analysis of functional brain imaging studies. Neurosci Biobehav Rev 42, 9–34 (2014).

39. Hill, C. A. et al. A causal account of the brain network computations underlying strategic social behavior. Nat Neurosci 20, 1142–1149 (2017).

40. Haruno, M. & Frith, C. D. Activity in the amygdala elicited by unfair divisions predicts social value orientation. Nat Neurosci 13, 160–161 (2010).

41. Carmichael, O. & Lockhart, S. Neurobiological programming of early life stress: Functional development of amygdala-prefrontal circuitry and vulnerability for stress-related psychopathology. Curr Top Behav Neurosci 289–320 (2018) doi:10.1007/7854.

42. Ginty, A. T., Kraynak, T. E., Kuan, D. C. & Gianaros, P. J. Ventromedial prefrontal cortex connectivity during and after psychological stress in women. Psychophysiology 56, 1–15 (2019).

43. Sinha, R., Lacadie, C. M., Todd Constable, R. & Seo, D. Dynamic neural activity during stress signals resilient coping. Proc Natl Acad Sci U S A 113, 8837–8842 (2016).

44. Burghy, C. a et al. Developmental pathways to amygdala-prefrontal function and internalizing symptoms in adolescence. Nat Neurosci 15, 1736–1741 (2013).

45. Phelps, E. A. & LeDoux, J. E. Contributions of the amygdala to emotion processing: From animal models to human behavior. Neuron vol. 48 (2005).

46. Maier, S. U., Makwana, A. B. & Hare, T. A. Acute Stress Impairs Self-Control in Goal-Directed Choice by Altering Multiple Functional Connections within the Brain’s Decision Circuits. Neuron 87, 621–631 (2015).

47. Philiastides, M. G., Biele, G. & Heekeren, H. R. A mechanistic account of value computation in the human brain. Proc Natl Acad Sci U S A 107, (2010).

48. Botvinick, M. M. Conflict monitoring and decision making: reconciling two perspectives on anterior cingulate function. Cogn Affect Behav Neurosci 7, 356–366 (2007).

49. Riccio, D. C., Ackil, J. & Burch-Vernon, A. Forgetting of stimulus attributes: Methodological implications for assessing associative phenomena. Psychol Bull (1992).

50. Bartz, J. A., Zaki, J., Bolger, N. & Ochsner, K. N. Social effects of oxytocin in humans: Context and person matter. Trends in Cognitive Science (2011).

51. Oosterbeek, H., Sloof, R. & Van De Kuilen, G. Cultural differences in ultimatum game experiments: Evidence from a meta-analysis. Exp Econ (2004).

52. Tzourio-Mazoyer, N. et al. Automated anatomical labeling of activations in SPM using a macroscopic anatomical parcellation of the MNI MRI single-subject brain. Neuroimage (2002) doi:10.1006/nimg.2001.0978.

53. Zhong, S., Chark, R., Hsu, M. & Chew, S. H. Computational substrates of social norm enforcement by unaffected third parties. Neuroimage 129, 95–104 (2016).

54. Hayes, A. F. Introduction to Mediation, Moderation, and Conditional Process Analysis. vol. 2507 (Guilford Press, 2017).

